# Multicellular Calcium Waves in Cancer-Associated Fibroblasts Regulate Neuronal Mimicry and Anisotropy Leading to Immune Exclusion

**DOI:** 10.1101/2025.10.23.684281

**Authors:** Giovanni Giangreco, Zoe Ramsden, Antonio Rullan, Steven Hooper, David Novo, Joana Leitão Castro, Probir Chakravarty, Marta Milan, Hamid Mohammadi, Sandra Llop, Marc Juarez, Deborah Schneider-Luftman, Marc Oliva, Laia Alemany, Mihaela Angelova, Aleksandra Olow, Xin Yu, Philip S Hobson, Kevin Harrington, Erik Sahai

## Abstract

Stromal barriers exclude CD8+ T cells from accessing cancer cells and hamper immune-mediated tumour control. Through multi-pronged analysis of tumours that transition from immune inflamed to immune excluded, we reveal that the formation of stromal barriers is associated with the acquisition of neuronal gene expression programmes in cancer-associated fibroblasts (CAFs), including TUBB3 expression. This leads to neuronal mimicry, with stromal barrier formation underpinned by coordinated transient bursts of intracellular calcium release, similar to those observed in neuronal tissue. Blockade of calcium release through either pharmacological or molecular interventions, such as STC2 depletion, prevents CAF alignment and the build-up of CD8+ T cells at stromal boundaries. Nintedanib treatment prevents neuronal mimicry and restores immune-mediated tumour control. Thus, we uncover unexpected mimicry of neuronal behaviour in CAFs, document the mechanism by which it leads to immune exclusion, and identify ways to prevent the induction of neuronal mimicry and restore immune-mediated tumour control.

**Graphical Abstract:** 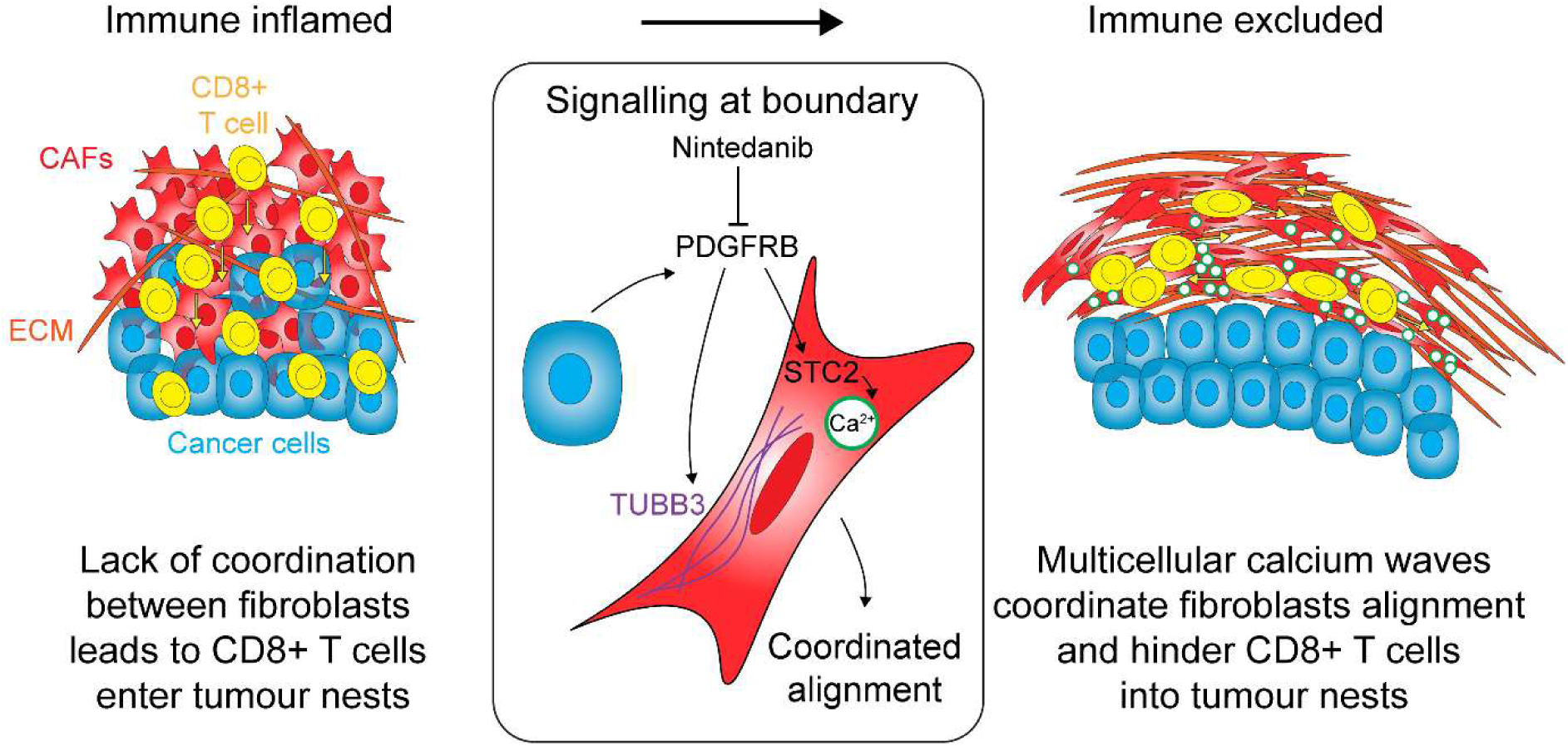

## Introduction

Tumours possess multiple mechanisms to evade immune surveillance [1]. These include interference with the ability of dendritic cells to activate CD8+ T cells, perturbation of the antigen presentation machinery in cancer cells, the expression of immune suppressive cytokines by cancer cells, and the recruitment of immune suppressive myeloid cells [1]. In addition, cancer associated fibroblasts (CAFs) can interfere with immune surveillance through the production of cytokines [2], [3], [4], [5], [6], [7], and by generating extracellular matrix (ECM) barriers that hinder CD8+ T cell access inside clusters of cancer cells, named tumour nests [8], [9], [10], [11], [12], [13], [14], [15]. These mechanisms can result in immune excluded tumours, where CD8+ T cells are present in stromal regions, but not in direct contact with tumour nests. Imaging studies have concluded that tumours can transition from an inflamed state to an immune excluded state [12], [16], [17], [18]; however, the molecular mechanisms underlying this transition are not well understood. Reversal of mechanisms underpinning immune exclusion is an appealing strategy for cancer therapies and is expected to enhance the efficacy of immune checkpoint blockade and improve the outcome of patients receiving immunotherapy.

CAFs are a highly abundant cell type present in the tumour microenvironment (TME) and key players in driving therapy resistance [19], [20], [21], [22]. CAFs can adopt diverse phenotypes, most notably myo-fibroblastic CAFs (myoCAFs), inflammatory CAFs (iCAFs) and antigen presenting CAFs (apCAFs) [23], a property observed also during development and wound healing [24]. myoCAFs are linked to Transforming Growth Factor Beta (TGFβ) activation, ECM deposition, and cancer cell metastasis. ECM deposition can be highly aligned anisotropic or more disorganised and isotropic, with the former pattern linked to more aggressive cancer phenotypes [25]. iCAFs are associated with immune cell recruitment / suppression, while apCAFs are connected to immune cell recruitment/activation [19], [23]. However, these categories are not rigid, nor strictly mutually exclusive; for example, LRRC15+ and MMP1+ myoCAFs have been associated with immune evasion through crosstalk with macrophages and CD4+ T regulatory cells respectively [26].

Extensive fibroblastic stroma is frequently associated with worse prognosis in squamous cell carcinoma (SCC) patients, in particular in head and neck SCC (HNSCC) [5]. HNSCC is responsible for about 1,000,000 cancer deaths annually [27]. Immunotherapy is effective against a subset of HNSCC, but most of them are not responsive despite generally having either a high tumour mutational burden or expressing viral oncoproteins [28], [29]. Numerous studies are seeking to improve immunotherapy responses; however, these are heavily focused on CD8+ T cell manipulations – targeting of LAG3, TIGIT, OX40 – with much less attention paid to modulation of stromal barriers that exclude CD8+ T cells.

In this study, we determine the mechanisms that promote immune exclusion and how to overcome them. Through a combination of highly multiplexed staining, live imaging and transcriptomics, we document the transition from immune inflamed to immune exclusion in murine SCC and human patients. We additionally establish a reductionist assay of tumour-stroma boundary that we exploit to uncover the molecular mechanisms by which CAFs promote immune exclusion. Both *in vivo* and *in vitro*, the interface between cancer cells and CAFs is characterised by dense arrays of CAFs aligned along the boundary that prevent CD8+ T cells accessing cancer cells. At the same time, CAFs acquire expression of neuronal marker – TUBB3 – and of neuro-inflammatory properties. Molecular analyses uncover a communication network active at this boundary that leads to the up-regulation of STC2 in CAFs and multicellular Ca^2+^ waves. Moreover, we show that repurposing of the anti-fibrotic drug, Nintedanib, prevents up-regulation of this network, neuronal mimicry, anisotropic organization, and restores immune-mediated tumour control.

## Results

### Mapping the transition from immune surveillance to immune escape in murine SCC

To study the interplay between the adaptive immune system and SCC, we used two syngeneic models known to represent carcinogen-induced oral SCC, MOC1 and MOC2 [30]. The two models exhibited different growth kinetics (Figure 1A). MOC1 tumours grew slowly between 5 and 15 days (doubling time ∼14 days), before accelerating to double every four days (Figure 1B). In contrast, MOC2 tumours grew exponentially (doubling time ∼3 days Figure 1B). The timing of the stasis of MOC1 tumour growth is consistent with an adaptive immune response, leading us to hypothesise that MOC1 tumours may be subject to immune surveillance around day 10. In support of this, MOC1 tumours grew rapidly with no period of stasis in Nod SCID gamma (NSG) mice (Figure S1A). To characterize the transition from immune surveillance to immune escape, we used single cell RNA sequencing (scRNAseq, Figure 1C&S1B). We interrogated the differences in the TME of MOC1 tumours at day 10 (stasis – referred to as MOC1 early) with MOC2 at day 10 (growth) and MOC1 around day 30 (growth – referred to as MOC1 late). Expression of antigen presenting machinery by MOC1 cells was mostly unchanged between early and late timepoints, suggesting that immune escape is not mediated through reduced antigen presentation (Figure 1D). Interestingly, MOC2 cells had lower expression of MHC class I related genes indicating that they might avoid immune surveillance due to defects in antigen presentation (Figure 1D). We next turned our attention to immune cells and noted an increase in immune exhaustion markers in CD8+ T cells in MOC1 late tumours (Figure 1E&S1C), consistent with a defective immune surveillance; this was not observed in MOC2 tumours. More holistic exploration of the data using CellChat analysis [31] revealed surprisingly few differences in communication involving lymphocytes between immune surveilled MOC1 early and immune escaped MOC1 late tumours (Figure 1F). Instead, the most prominent difference in CellChat analysis was altered cancer cell – stromal crosstalk (Figure 1F). These data prompted us to investigate changes in CAFs as MOC1 tumours overcome immune surveillance. Intriguingly, we found no clear change in the relative abundance of different sub-types across samples (Figure S1D&E), suggesting that CAFs were not switching between previously established sub-types. We therefore interrogated cancer cell – CAF crosstalk in MOC1 late tumours in more depth. This implicated multiple receptor tyrosine kinase ligands signalling from cancer cells to CAFs, and different ECM components potentially mediating bi-directional signalling (Figure S1F). Further analysis of the ligands and receptors involved in this crosstalk revealed a surprising enrichment for genes linked to neuronal state, hinting a possible change in CAF functionality, migration and substrate adhesion (Figure 1G). Moreover, we observed an enrichment for neuroinflammatory signatures in the CAF dataset in immune escaped MOC1 late tumours (Figure 1H). Figure S1G shows that multiple neuronal associated genes exhibited increased expression in MOC1 late tumours compared to MOC1 early, with particularly high expression observed for the neuronal marker *Tubb3* [32] (Figure 1I).

**Figure 1:**
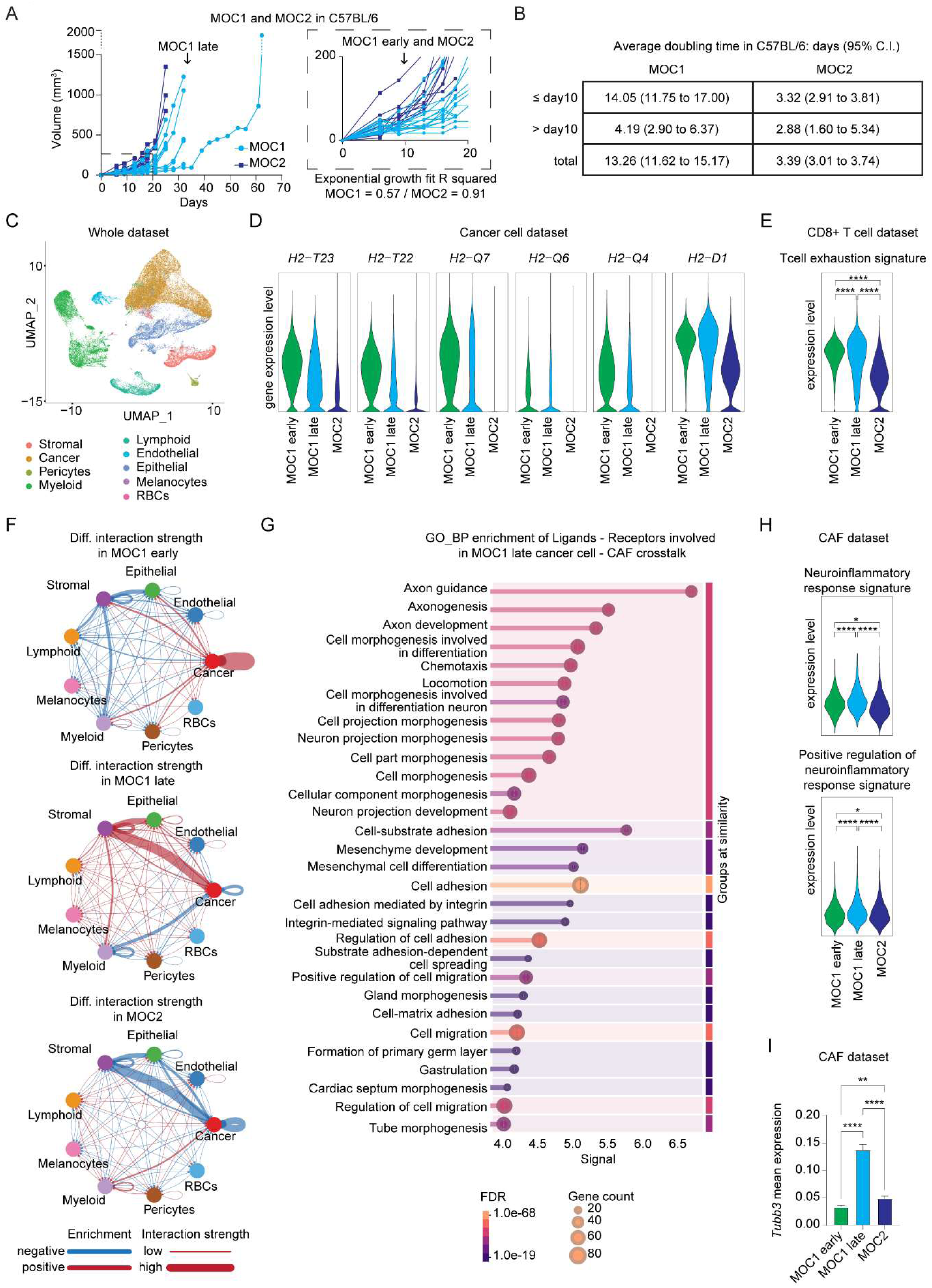
CAF – cancer cell crosstalk is boosted after immune evasion via acquisition of CAF-specific neuronal features. (A) Growth curve of sub-cutaneous injection of MOC1 and MOC2 in C57BL/6 mice. Black dashed box highlights growth pattern difference around day10. Arrows indicate the selected experimental time points for MOC1 early, MOC1 late and MOC2. R squared of exponential growth is shown. N=15 mice (MOC1) and N=10 mice (MOC2). (B) Average doubling time in days (with 95% confidence interval) for mice plotted in Figure 1A from exponential growth fit calculation. (C) Uniform Manifold Approximation and Projection (UMAP) of MOC1 early, MOC1 late and MOC2 scRNAseq. N=3 mice per group. (D) Violin plot of selected genes. (E) Violin plot of the indicated signature from [66]. P value from two-sided Wilcoxon test. (F) Circle plot showing the difference in CellChat interaction strength in the indicated samples. For each sample (i.e. MOC1 early) the comparative analysis was performed against the other samples (i.e. MOC1 late and MOC2). The weight of edges represents the difference in interaction strength. The colour represents the direction of change relative to the comparison. Red = increased in the specified sample; Blue = decreased in in the specified sample. (G) Multivariable plot showing enrichment of top 30 Gene Ontology Biological Processes (GO_BP) signatures associated with cancer cell – CAF crosstalk in MOC1 late sample. X axis shows STRING signal values. Bubble size represents gene counts. Bar colour on the right represents FDR. Signatures are grouped by similarity (threshold 0.8). Plot generated with String web portal [67]. (H) Violin plot of the indicated signatures. P value from two-sided Wilcoxon test. (I) Box plot of *Tubb3* mean expression in CAF cell population from scRNAseq. Average + SEM are shown. One-way ANOVA Holm-Sidak corrected for multiple comparison.

To characterize further the TME in MOC1 early, MOC1 late and MOC2 and investigate the presence of CAF-related neuronal features, we performed imaging mass cytometry (IMC) of the same samples. We used an antibody panel that includes markers for the well-established TME cellular populations and ECM components (Figure S2A), with TUBB3 added as marker linked to neuronal state. Following staining and image acquisition, we performed cellular segmentation (Figure 2A) and classified cells according to known cellular lineages [33] (Figure S2B). Figures 2B&C show representative images and the relative abundance of different cell types in the MOC1 early, MOC1 late, and MOC2 tumours, respectively. MOC1 early tumours had higher levels of CD8+ T cells, CD4+ T cells, and dendritic cells compared to MOC1 late and MOC2 tumours, which is consistent with inflamed immune TME in MOC1 early tumours. Both MOC1 early and late tumours had more CAFs than MOC2, and, linking to the transcriptomic analysis, TUBB3 was expressed in a higher proportion of CAFs in MOC1 late tumours (Figure 2D&E).

**Figure 2:**
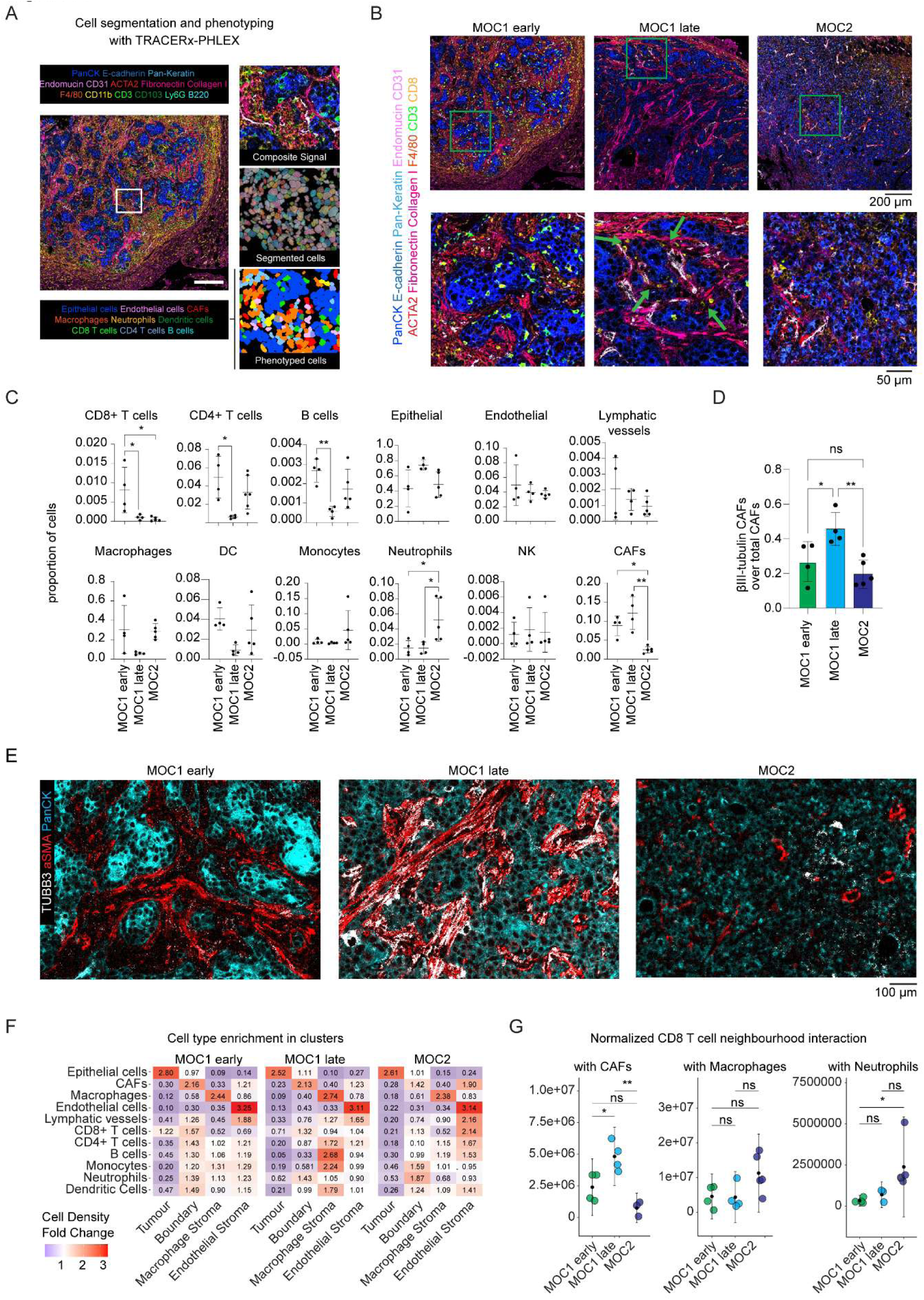
CD8+ T cells become excluded in MOC1 late tumour and are stacked at tumour-stroma boundary. (A) Cell segmentation and phenotyping was performed using the TRACERx-PHLEX [33] pipeline for highly multiplexed image analysis. Representative schematic showing composite signal, segmented cell objects and phenotyped cells in a MOC1 early tumour. (B) Representative images of MOC1 early, MOC1 late and MOC2. Green arrows indicate examples of CD8+ T cells localized at tumour-stroma boundary in MOC1 late. (C) Box plot with mean ± standard deviation (SD) and dots of cellular proportion of each annotated cell type in MOC1 early (N=4), MOC1 late (N=4) and MOC2 (N=5). One-way ANOVA Tukey corrected for multiple comparison. (D) Box plot showing proportion of TUBB3 positive CAFs over total CAFs in MOC1 early (N=4), MOC1 late (N=4) and MOC2 (N=5). One-way ANOVA Holm-Sidak corrected for multiple comparison. (E) Exemplar images of TUBB3, ACTA2 and pan cytokeratin (PanCK). (F) Heatmap of fold change of proportional cell density in K = 4 clusters identified in unsupervised clustering, calculated per tissue type (N = 13). (G) Frequency of CAFs, macrophages and neutrophils present in the neighbourhood of CD8+ T cells across MOC tissues, normalised to overall CD8+ T cell density per tissue. N = 13, Two-tailed unpaired Student’s t test, p values Benjamini-Hochberg adjusted.

To gain insight into the spatial organisation of the tumours and inter-cellular interactions, we performed neighbourhood analysis (Figure S2C). K-means clustering indicated four predominant neighbourhoods (Figure S2C), which corresponded to tumour nests, CAF boundary, macrophage stroma and peri-vascular stroma (Figure S2D). Figure 2F shows the abundance of different cell types in these four regions. Of note, CD8+ T cells were enriched in tumour nests in MOC1 early tumours, but not in MOC1 late tumours. Similar results were obtained when tumours were binarized into cancer cell or stroma regions (Figure S2E&F). The shift of CD8+ T cells away from tumour cells was accompanied by an increase in CAFs in the same neighbourhood as CD8+ T cells (Figure 2G and green arrows in Figure 2B). In contrast, the association of CD8+ T cells with macrophages or neutrophils was unchanged between MOC1 early and late tumours (Figure 2G). MOC2 tumours had both globally elevated neutrophils and increased neutrophils in CD8+ T cell neighbourhoods (Figure 2C&F), once again suggesting a different mechanism of immune evasion. Together, these data indicate that MOC1 tumours transition from an initial slow growing immune inflamed state to an exponentially growing immune excluded one, with increased CD8+ T cell – CAF physical interactions and communication in immune excluded tumours with acquisition of neuronal features from CAFs.

### Impaired CD8+ T cell migration in immune excluded tumours is guided by CAFs and ECM

Cell migration underpins the distribution of CD8+ T cells; therefore, we performed live imaging of CD8+ T cells in tissue slices of MOC1 early and MOC1 late tumours grafted into ACTA2 (ACTA2)::DsRed reporter mice to visualise CAFs (Figure 3A&S3A). Figure S3B shows representative images of the MOC1 early and MOC1 late tumours. As expected, fewer CD8+ T cells are observed in tumour nests in MOC1 late tumours (Figure S3C). Cell tracking revealed that the migration of CD8+ T cells in MOC1 late tumours was markedly slower (Figure S3D and Movie 1&2), and the few CD8+ T cells that were observed in MOC1 late tumour nests had further reduced velocity (Figure 3B&S3E and Movie 3). Moreover, CD8+ T cells at tumour-stroma boundaries were frequently observed migrating parallel to stroma boundaries and rarely crossed from the stroma into the tumour nests, suggesting that some aspect of the tumour – stroma boundary may guide CD8+ T cells to move around the tumour nest, but not enter it (Figure 3C and Movie 4). Indeed, we noticed that the major axis of CD8+ T cells was parallel to CAF alignment (green arrows in Figure 3C and Movie 4). This motivated us to consider the spatial organisation and alignment of the ECM and CAFs, and how they related to the tumour-stroma boundary. These features were delineated using the TWOMBLI algorithm [34], designed for the detection and analysis of fibres (Figure 3D). Analysis of Collagen I and Fibronectin 1 (FN1) fibres and ACTA2-positive CAFs indicated that they were less curved and more aligned in MOC1 late tumours compared to MOC1 early and MOC2 (Figure 3E&S3F). The orientation of alignment was parallel to the tumour-stroma interface in MOC1 tumours (Figure 3F&S3G). Besides displaying lower stromal alignment, MOC2 tumours exhibited poorly defined tumour-stroma boundaries, with individual cancer cells frequently invading into the stroma (Figure 3E&3F&S3F&S3G).

**Figure 3:**
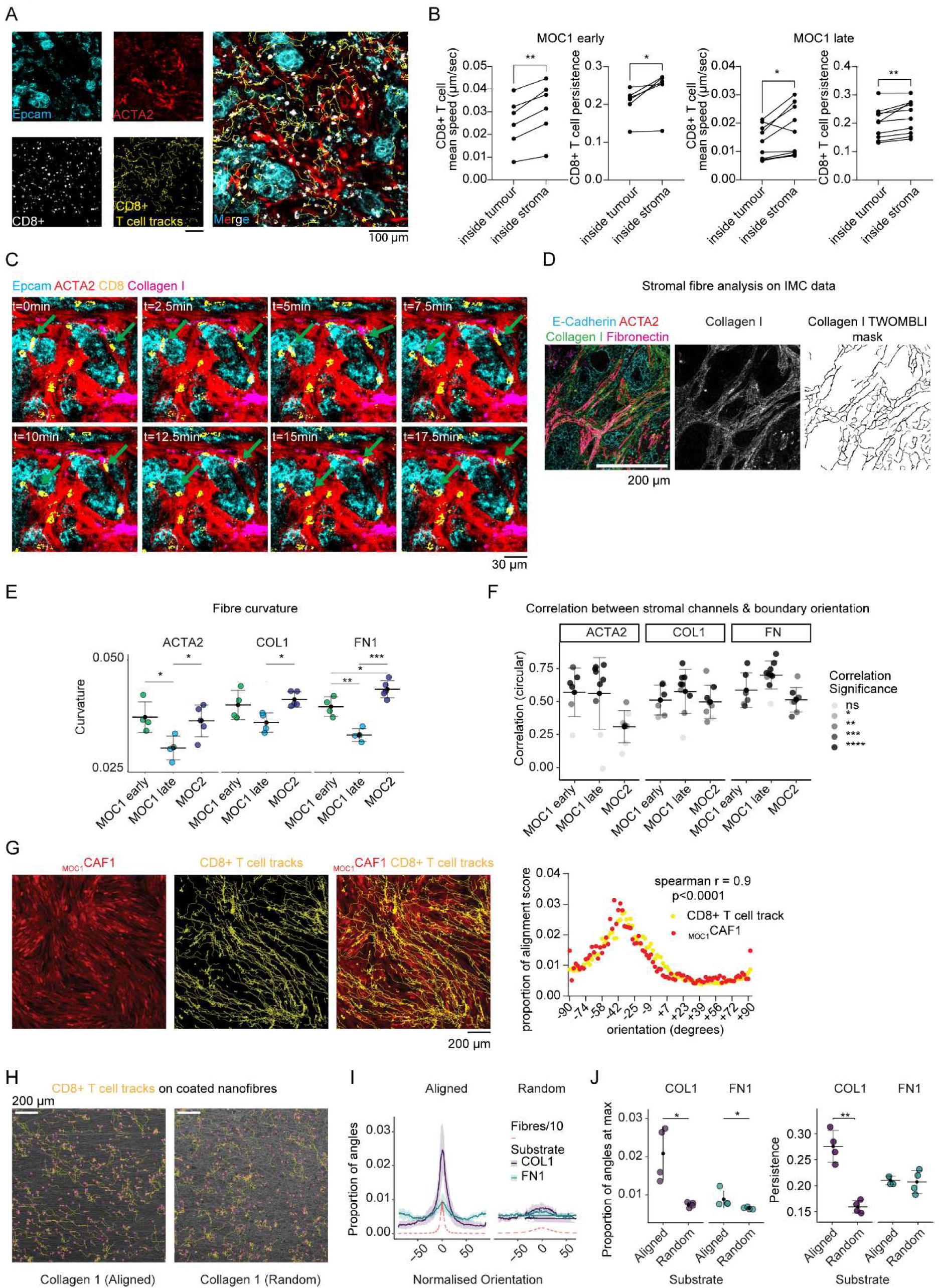
CD8+ T cell migration is directed by CAF alignment. (A) Exemplar z-stack projection of a MOC1 tissue slices stained with CD8+ T cell and Epcam (for MOC1 cancer cells), ACTA2 (CAFs) and CD8+ T cell tracks. (B) Dot plot of CD8+ T cell track metrics separated by tumour or stroma in MOC1 early (N=6 per group) and MOC1 late (N=10 per group). Two-tailed paired Student’s t test. (C) Exemplar z-stack image series of CD8+ T cells, Collagen I, Epcam (for MOC1 cancer cells), ACTA2 (CAFs). Green arrows show 2 different CD8+ T cells with elongated morphology moving around the nest following CAFs and ECM alignment. (D) Schematic of TWOMBLI pipeline to extract fibre masks from IMC stromal antibody staining. (E) TWOMBLI metrics for fibre masks from stromal channels, averaged per sample. N = 13, Student’s unpaired Student’s t test, p values Benjamini-Hochberg adjusted. (F) Circular correlation between orientation of tumour boundary and stromal fibres in MOC tissues. Alpha scale represents significance of correlation per image (N = 23 images). (G) Left, example image of _MOC1_CAF1 spatial organization and CD8+ T cell tracks spatial distribution. Right, XY axis plot of orientation angle and proportion of alignment score for CD8+ T cell track and _MOC1_CAF1 major axis. Sperman r correlation metric is shown. (H) Representative images of primary CD8+ T cells migrating on collagen-coated nanofibres. Yellow tracks indicate CD8+ T cell trajectory. (I) Proportional distribution of orientations of CD8+ T cell tracks and nanofibres calculated, normalised to major fibre angle = 0. CD8+ T cell track distribution plotted as an average of N=4 experiments ± SD. Fibre distribution plotted as an average of all experiments and shown at a scale of 1 x 10e^-1^. (J) Left, proportion of total CD8+ T cell track angles at peak of orientation distribution. N=4. Unpaired Wilcoxon test, p values Benjamini-Hochberg adjusted. Right, CD8+ T cell migration persistence on different ECM substrates and topologies. N=4. Two-tailed unpaired Student’s t test, p values Benjamini-Hochberg adjusted.

In order to test whether our observations occur also in human samples, we characterized a cohort of oropharynx HNSCC HPV negative patients by staining for markers of epithelial cancer cells (E-Cadherin and Pan-Cytokeratin), CAFs (ACTA2), ECM (Collagen 1), CD8+ T cells and we also included TUBB3. Of note, we frequently found TUBB3 positive CAFs (Figure S4A). We observed different patterns of CD8+ T cell distribution, with some examples where CD8+ T cells were found infiltrate in tumour nests (yellow arrows in Figure S4A), while others where they were physically excluded from nests (green arrows in Figure S4A). Importantly, we observed that CD8+ T cells excluded often showed the length of major axis being parallel to CAF alignment (green arrows in Figure S4A) and overlapping with TUBB3 positive fibroblasts. Overall, these data indicate that our observations with mouse models are observed also in patients’ samples.

To investigate whether CAFs were guiding CD8+ T cell migration parallel to the boundary, thereby stopping them to enter tumour nests, we isolated CAFs from MOC1 late tumours (hereafter called _MOC1_CAF, Figure S4B). We isolated also CAFs from MOC2 tumours (hereafter called _MOC2_CAF) for comparison. Interestingly, the *in vitro* alignment and ECM deposition of the isolated CAFs matched their *in vivo* phenotypes, with _MOC1_CAFs generating aligned anisotropic patterns and _MOC2_CAFs generating isotropic patterns (Figure S4C-E). Imaging of CD8+ T cells co-cultured with _MOC1_CAF demonstrated that _MOC1_CAFs promoted CD8+ T cell migration in the direction of CAF alignment (Figure 3G and Movie 5). CellChat analysis showed that Collagen was the most likely mechanism for CAF – CD8+ T cell interaction (Figure S4F). To test if the ECM could be sufficient to guide CD8+ T cells, we coated either anisotropic aligned or isotropic non-aligned arrays of nanofibres with Collagen I or FN1 and tracked T cell migration (Figure 3H). Parallel Collagen I and, to a lesser extent, FN1 were able to guide CD8+ T cell migration (Figure 3I&J), leading to more persistent migration (Figure 3J). Cell speed and displacement were not affected by alignment (Figure S4G). These data suggest that CAF alignment and ECM guide CD8+ T cell migration parallel to the boundary, preventing them to enter tumour nests in immune excluded tumours.

### CAF alignment correlates with CD8+ T cell accumulation at reconstituted tumour-stroma boundaries

To explore CAF-guided CD8+ T cell migration, we established a tri-culture assay replicating a tumour-stroma boundary. Cancer cells and CAFs were plated on either side of a polydimethylsiloxane (PDMS) insert. When cells were confluent, the insert was removed and cancer cells migrated until they were in contact with CAFs forming a boundary, then activated mouse primary CD8+ T cells were added (Figure 4A). Concordant with our *in vivo* observations, MOC1 cancer cells and _MOC1_CAFs generated a well-defined linear boundary, whereas MOC2 and _MOC2_CAF generated a poorly defined boundary with MOC2 cells infiltrating the stromal area (Figure S5A to be compared with Figure 2B&F). We also noticed a higher density of CAFs at MOC1/_MOC1_CAF boundaries, but not MOC2/_MOC2_CAF boundaries (Figure S5A). The organisation of actomyosin cables and FN1 fibres is parallel to the boundary in the MOC1 assays (Figure 4B). Combinations of MOC1 with _MOC2_CAF and vice versa yielded intermediate phenotypes (Figure S5A).

**Figure 4:**
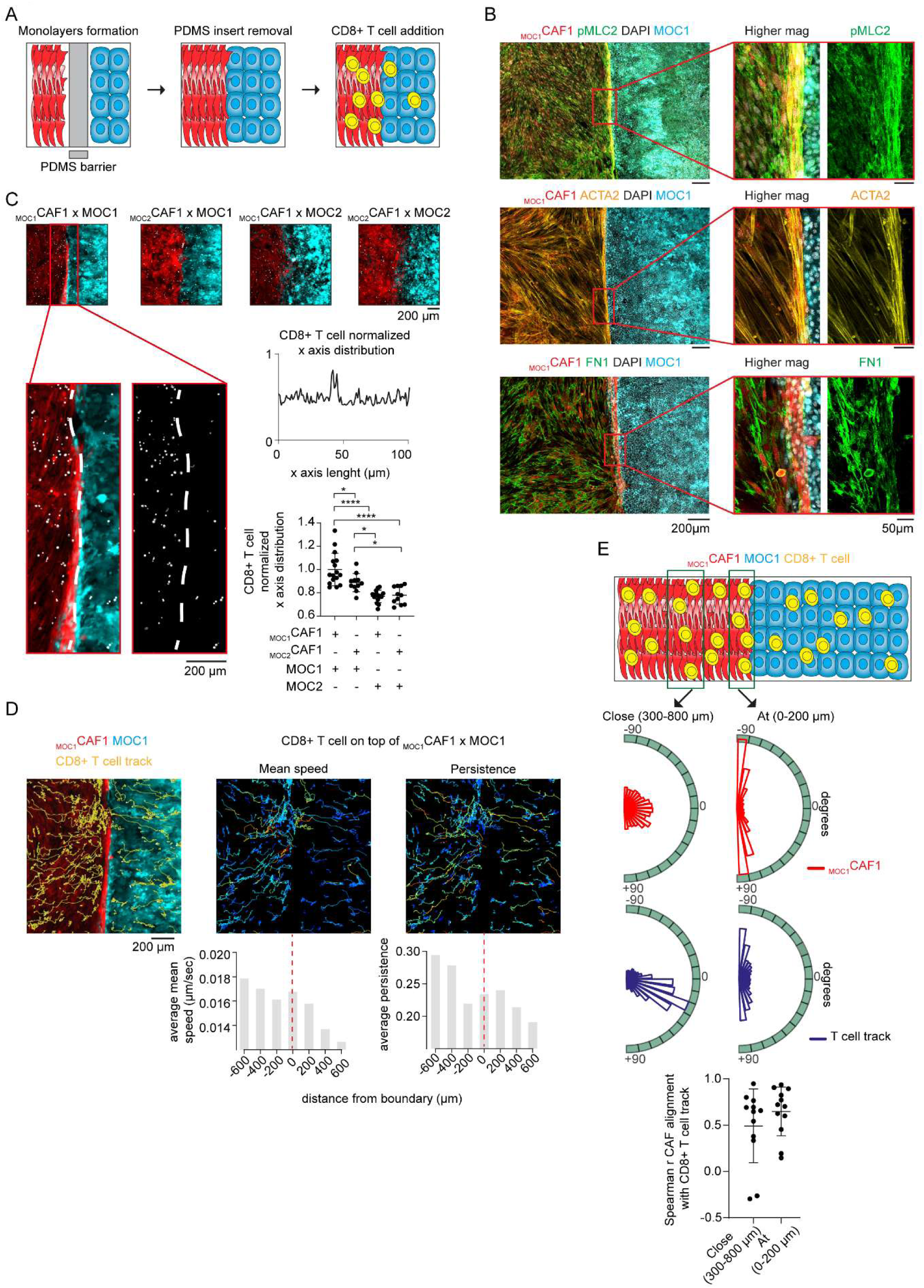
CAF alignment at reconstituted tumour-stroma boundaries prevents CD8+ T cell infiltration. (A) Schematics of tumour-stroma reconstitution assay with CD8+ T cell addition. (B) Exemplar images of MOC1-_MOC1_CAF staining at reconstituted tumour-stroma boundary. (C) Top, exemplar images of reconstituted tumour-stroma boundaries with CD8+ T cell addition. Middle right, example of a normalized CD8+ T cell x axis distribution. Bottom right, dot plot with mean ± SD of CD8+ T cell normalized x axis distribution. N≥11. One-way ANOVA Holm-Sidak corrected for multiple comparison. (D) Top, schematics of the assay. Middle, exemplar images of reconstituted tumour-stroma boundaries with MOC1-_MOC1_CAF with CD8+T cell tracks. Colour coded T cell tracks represent CD8+ T cell mean speed (centre) and persistence (right). Bottom, box plot plotting average CD8+ T cell metrics as function of distance from boundary. (E) Top, Circular plot showing orientation of CAF alignment (red) and T cell track (blue) at two different locations from the boundary. Bottom, Dot plot showing Spearman r correlation of CAF orientation and CD8+ T cell track orientation. N=12.

We next monitored CD8+ T cell migration and distribution along these boundaries. Figure 4C shows that CD8+ T cells accumulate at the boundary of MOC1/_MOC1_CAF cultures, but not MOC2/_MOC2_CAF cultures. This behaviour was quantified by plotting the normalized distribution of CD8+ T cells on the x-axis, with a clear peak in CD8+ T cell density at the boundary observed in MOC1/_MOC1_CAF cultures (Figure 4C). Timelapse imaging revealed that CD8+ T cells in the stromal region migrate towards the boundary, but rarely cross (Figure 4D and Movie 6). Similar to our observations in tissue slice cultures (Figure 3B), CD8+ T cell migration is slower in cancer cell regions compared to CAF regions, (Figure 4D&S5B). Upon approaching the boundary, CD8+ T cells migrate in parallel to the boundary, with a high correlation of migration track angles and CAF orientation (Figure 4E). These observations were replicated using a human SCC model system composed of A431 SCC cells, VCAF2b (anisotropic SCC-derived CAFs), and primary human CD8+ T cells (Figure S6A). As in the murine system, CAFs aligned parallel to the boundary and the migration of CD8+ T cells along the CAFs lead to their accumulation at the boundary (Figure S6B-D). These analyses demonstrate that key features of CD8+ T cell behaviour at tumour-stroma boundaries can be replicated *in vitro* and that CAF alignment at the boundary is a likely mechanism preventing CD8+ T cell crossing.

### Nintedanib enables CD8+ T cell access into tumour nests and improves tumour control

We next sought to use our assay to identify pathways regulating the migration of CD8+ T cells, with the goal to find mechanisms to disturb their accumulation at the tumour-stroma boundary and increase penetration into tumour nests. A focused array of pharmacological inhibitors known to impact CAF biology were tested (Figure 5A). The strongest effect of CD8+ T cell accumulation was observed with nintedanib, which is a kinase inhibitor used for the treatment of lung fibrosis [35] and known to target Platelet-Derived Growth Factor Receptor (PDGFR), Vascular-Endothelial Growth Factor Receptor (VEGFR) and Fibroblast Growth Factor Receptor (FGFR) (Figure 5A&B). Following nintedanib treatment, CAFs at the boundary were less densely packed and not as strongly aligned (Figure 5B&C). Nintedanib also reduced the alignment of _MOC1_CAF in confluent mono-culture (Figure S7A). Nintedanib had little effect on cell proliferation at the concentration used for migration assays (1 µM) (Figure S7B). We confirmed that nintedanib affects CAF alignment in three other human CAF models known to be strongly anisotropic [25] (Figure S7C). Timelapse imaging revealed that nintedanib reduced CD8+ T cell migration parallel to the tumour-stroma boundary and increased the proportion of crossing cells (Figure 5D and Movie 7&8). Intriguingly, nintedanib reduced expression of the neuronal marker TUBB3 in _MOC1_CAF1, suggesting a possible connection between neuronal gene expression and anisotropic organisation of CAFs (Figure 5E).

**Figure 5:**
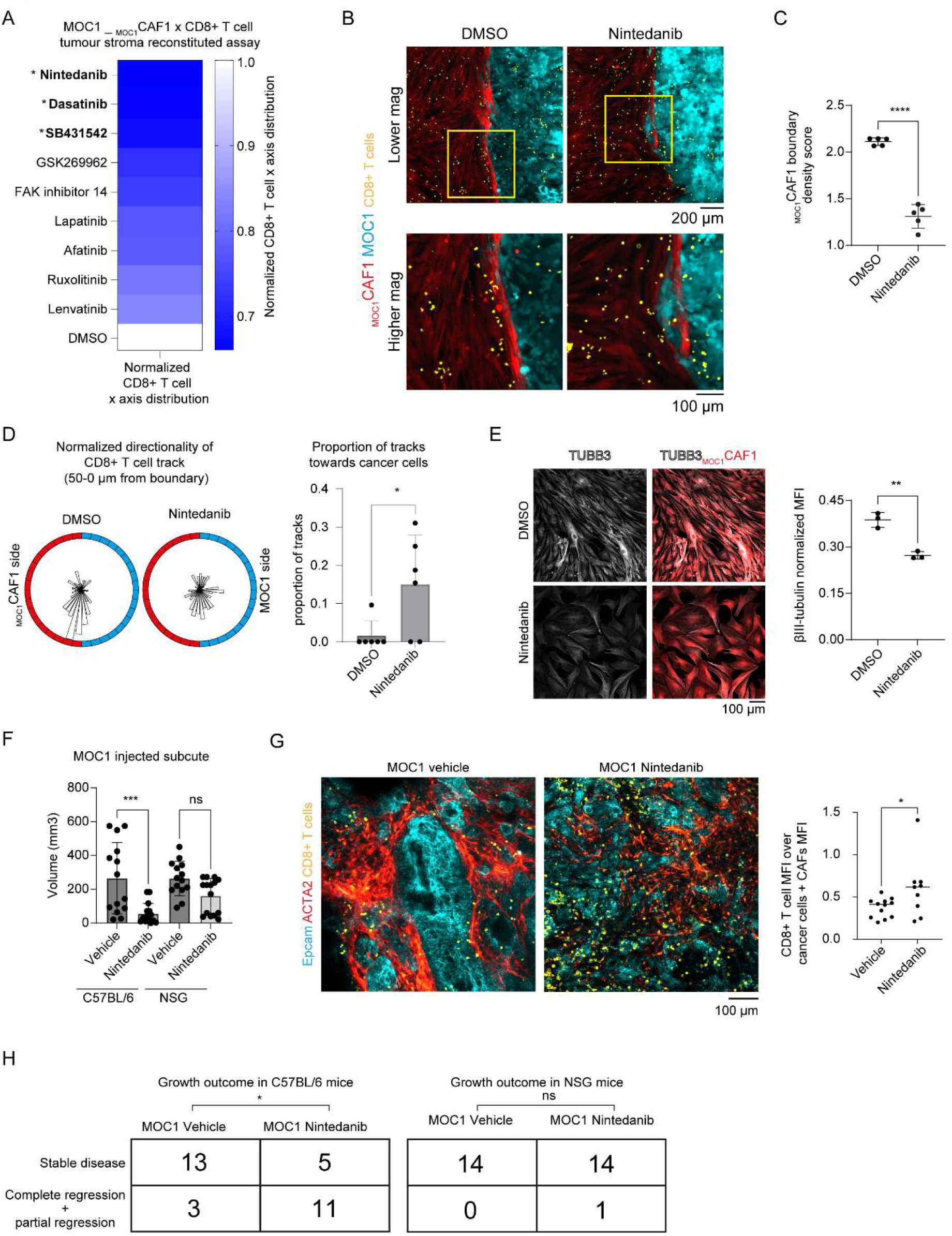
Nintedanib treatment alters tumour-stroma boundary architecture, increases CD8+ T cell infiltration and improves tumour control. (A) Heatmap of CD8+ T cell normalized distribution over x axis in the reconstituted tumour-stroma boundary assay with MOC1 and _MOC1_CAF1 with the indicated treatments. N≥3. Brown-Forsythe and Welch ANOVA Benjamini corrected for multiple comparison (*, p value < 0.5). (B) Exemplar images of tumour-stroma boundary with addition of CD8+ T cells in DMSO control condition and with Nintedanib 1 μM treatment. (C) Dot plot with mean ± SD of boundary density score for the indicated conditions. N=5. Two-tailed unpaired Student’s t test. (D) Left, circular plot of CD8+ T cell track orientation distant up to 50 μm from the boundary from the CAF side. Right, box plot with mean + SD and dots of proportion of CD8+ T cell tracks towards boundary in the indicated conditions. N=6. Two-tailed unpaired Student’s t test. (E) Left, exemplar images of TUBB3 expression in indicated samples in _MOC1_CAF1. Right, dot plot with mean ± SD of TUBB3 normalized MFI. N=3. Two-tailed unpaired Student’s t test. (F) Box plot and dots with mean ± SD of tumour volume at 26 days from sub-cutaneous injection of MOC1 in C57BL/6 or NSG mice. N≥14. One-way ANOVA Holm-Sidak corrected for multiple comparison. (G) Left, exemplar images of CD8+ T cells, Epcam (for cancer cells), ACTA2 (CAFs) for the indicated conditions. Right, dot plot with mean of CD8 mean fluorescent intensity (MFI) over sum of Epcam and DsRed MFI in MOC1 vehicle (N=12) and MOC1 Nintedanib (N=9). Two-tailed unpaired Student’s t test. (H) Table of growth outcome in C57BL/6 and NSG mice treated with vehicle control or nintedanib. Two-sided Fisher’s exact test.

These data indicate that nintedanib affects CAF alignment and organization at the boundary, increasing access of CD8 T+ cells into tumour nests and thereby increasing tumour control. This hypothesis was confirmed *in vivo* by treating mice bearing MOC1 with nintedanib, which led to a significant reduced tumour burden in immuno-competent mice (Figure 5F&S7D). We observed increased number of CD8+ T cells in tumour nests of MOC1 tumours (Figure 5G). Of note, Nintedanib had a minimal effect of MOC1 tumours in immuno-deficient mice (Figure 5F&5H&S7E). Moreover, MOC2 tumours did not respond to nintedanib treatment (Figure S7F). Overall, these data indicate that nintedanib targets nests-like tumour and requires a functional immune system for effective control.

### Identification of a regulatory network controlling CD8+ T cell accumulation at stromal boundaries

To obtain more specific molecular insights into MOC1 – _MOC1_CAF crosstalk occurring at tumour-stroma boundaries and the mechanism by which nintedanib might act, we performed bulk mRNAseq on MOC1/_MOC1_CAF1 co-cultures and mono-cultures treated with either DMSO or nintedanib (Figure 6A). Transcriptomic analysis using single-sample Gene Set Enrichment Analysis (ssGSEA) revealed several interesting sets of genes whose expression in CAFs was modulated by cancer cells in a nintedanib-dependent manner (Figure S8A). In particular, we noted that _MOC1_CAF1 showed upregulation of processes linked to neuronal development, actomyosin contractility, PKC regulation, protein folding and endoplasmic reticulum (ER) biology (Figure 6B with exemplar gene sets shown in Figure S8B). Consistent with Figure 5E, *Tubb3* mRNA expression was upregulated in CAFs co-cultured with MOC1 and downregulated when nintedanib was present (Figure 6C). In line with expectation, we observed up-regulation of TGFβ and TNFα pathways in co-cultured _MOC1_CAF1, but this was not impacted by nintedanib (Figure S8A).

**Figure 6:**
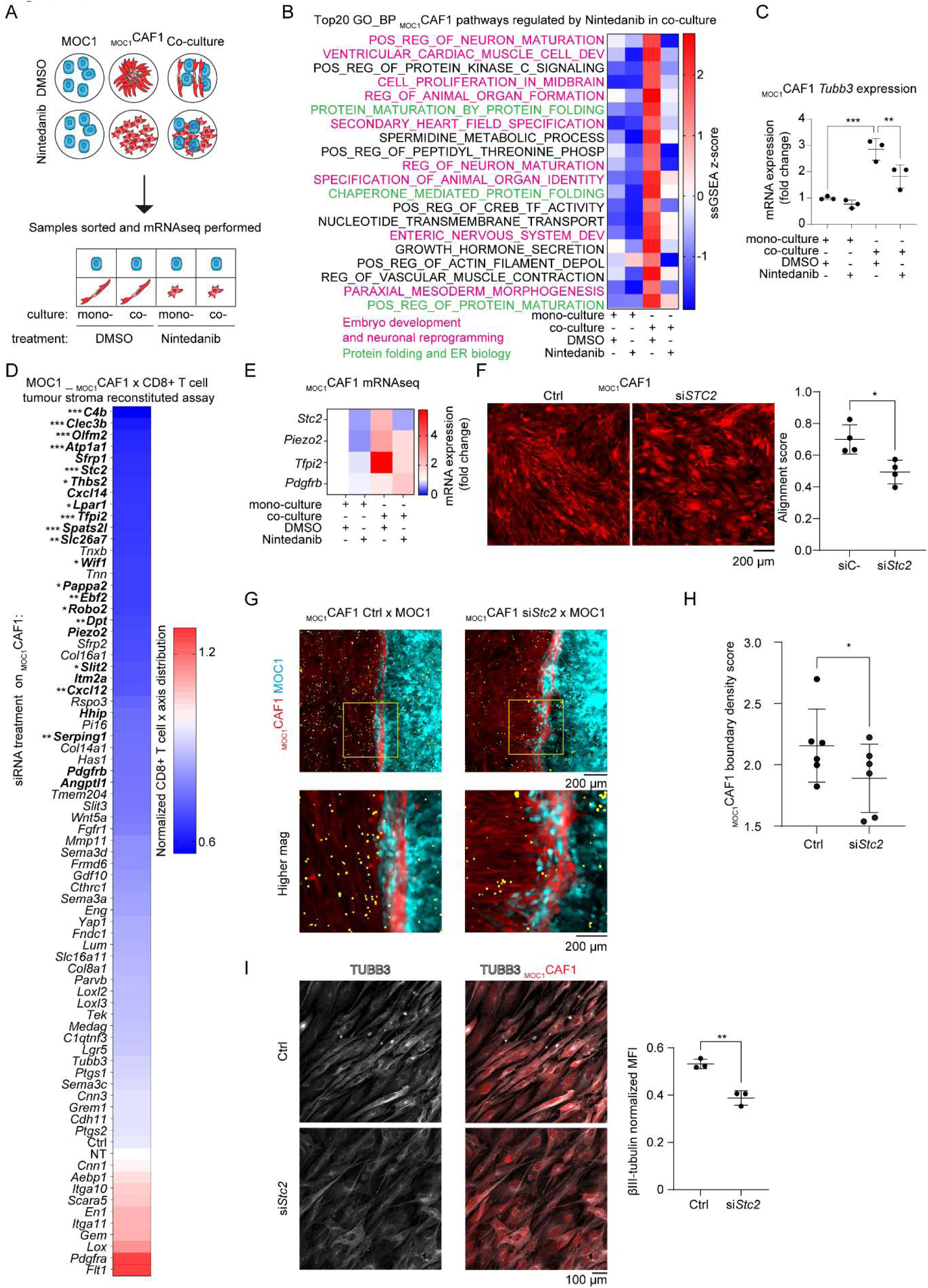
STC2 expression at tumour-stroma boundaries is upregulated in CAFs, dictating alignment and CD8+ T cell spatial distribution. (A) Schematics of MOC1 – _MOC1_CAF co-culture, treatment, sorting and mRNAseq strategy. (B) Heatmap of ssGSEA z-score of top20 gene ontology biological processes (GO_BP) pathways regulated by Nintedanib in co-culture in _MOC1_CAF. (C) Dot plot with mean ± SD of *Tubb3* mRNA expression in the mRNAseq dataset for the indicated conditions. N=3. Paired one-way ANOVA Holm-Sidak corrected for multiple comparison. (D) Heatmap of CD8+ T cell normalized distribution over x axis in the reconstituted tumour-stroma boundary assay with MOC1 and _MOC1_CAF1. The gene list represents siRNA treatment performed on _MOC1_CAF1. N≥3. Brown-Forsythe and Welch ANOVA Benjamini corrected for multiple comparison (***, p value < 0.001; **, p value < 0.01; *, p value < 0.5, bold, p value <0.1). (E) Heatmap of gene expression of the selected genes of interest. (F) Left, exemplar images of _MOC1_CAF1 for the indicated conditions. Right, dot plot with mean ± SD of CAF alignment score. N=4. Two-tailed unpaired Student’s t test. (G) Exemplar images of tumour-stroma boundary and addition of CD8+ T cells with _MOC1_CAF1 siC- or si*Stc2*. (H) Dot plot with mean ± SD of boundary density score for the indicated conditions. N=6. Two-tailed paired Student’s t test. (I) Left, exemplar images of TUBB3 expression in indicated samples. Right, dot plot with mean ± SD of TUBB3 normalized MFI. N=3. Two-tailed unpaired Student’s t test.

To determine the functional relevance of up-regulated genes to CD8+ T cell accumulation at tumour boundaries, we selected 73 genes based on their expression in the scRNAseq CAF dataset, bulk mRNAseq and a published dataset of isotropic / anisotropic fibroblasts [25] (Figure S9A). Figure 6D shows the results of siRNA-mediated depletion of these genes in _MOC1_CAF on the accumulation of CD8+ T cells at the boundary. Silencing of kinase known to be inhibited by nintedanib [35] showed that only *Pdgfrb* downregulation decreased CD8+ T cell accumulation, while *Pdgfra*, *Fgfr1* or *Flt1* did not (Figure 6D). Multiple Nintedanib-regulated genes were required for CD8+ T cell accumulation in the boundary zone, including *Pdgfrb*, *Stc2*, *Piezo2*, and *Tfpi2* (Figure 6E). However, *Tubb3* knock-down did not show an effect on CD8+ T cell boundary accumulation, indicating that its expression is a downstream marker of the neuronal state but does not have a functional role in this context. We focused on Stc2 (Stanniocalcin-2) as it was among the top-ranked genes identified (Figure S8C) and it is reported to play a role in neuronal differentiation and regeneration [36], [37], and is linked to receptor tyrosine kinase signalling [38], [39]. *Stc2* depletion (Figure S9B) reduces CAF alignment in _MOC1_CAF1 confluent monolayer (Figure 6F), reduces CAF density at the tumour-stroma boundary and CD8+ T cell accumulation (Figure 6G&H), phenocopying nintedanib treatment. *Stc2* depletion also reduced TUBB3 expression in _MOC1_CAF1, suggesting that it is functionally important for the acquisition of neuro-like features in CAFs (Figure 6I). We validated the effect of *STC2* depletion on CAF alignment with 3 other anisotropic human CAF cell lines (Figure S9C). Of note, *STC2* expression correlates with the CAF markers *ACTA2* and *PDGFRB, and* worse overall survival and progression free survival in HPV negative HNSCC (Figure S9D&E). These data favour a model in which tumour-stroma boundary boosts a regulatory network with increased expression of PDGFRB, leading to elevated expression of STC2 and subsequently the expression of neuronal markers, CAF organisation parallel to tumour-stroma boundaries and the diversion of CD8+ T cell migration.

### CAF alignment at the boundary boosts multicellular calcium waves

We explored how the signalling described above might regulate CAF alignment at tumour-stroma boundaries, and, more specifically, if the changes in neuronal-linked genes might have important functional consequences. Coordinated multicellular pulses of Ca^2+^ are a cardinal feature of neurons, while both STC2 and nintedanib have been reported to regulate intracellular Ca^2+^ levels [40], [41]. Moreover, we observed nintedanib-dependent upregulation of calcium-related gene signatures in _MOC1_CAF1 co-cultured with MOC1 (Figure 7A). Therefore, we sought to image Ca^2+^ dynamics to determine its contribution to cancer cell and CAF crosstalk. _MOC1_CAF expressing the Ca^2+^ sensor GCaMP6 were generated and used in the *in vitro* boundary assay (Figure 7B and Movie 9). First, we performed spike detection at the pixel level (Figure 7C) and characterized properties associated with intensity and duration (Figure S10A). Then, we performed spatiotemporal clustering of the spikes to analyse their properties as function of distance from the boundary (Figure 7D). Strikingly, coordinated multicellular calcium waves (MCWs) were observed and became more intense (Figure 7E&F), more frequent in space (Figure 7E&S10B), bigger in size (Figure S10C&D) and shorter in duration (Figure S10E&F) the closer they were to the boundary. Crucially, treatment with nintedanib and depletion of *Stc2* both reduced MCW intensity at the boundary (Figure 7G-J). Overall, these data show that CAFs at tumour-stroma boundaries upregulate MCWs and that either nintedanib or *Stc2* depletion are sufficient to decrease their intensity.

**Figure 7:**
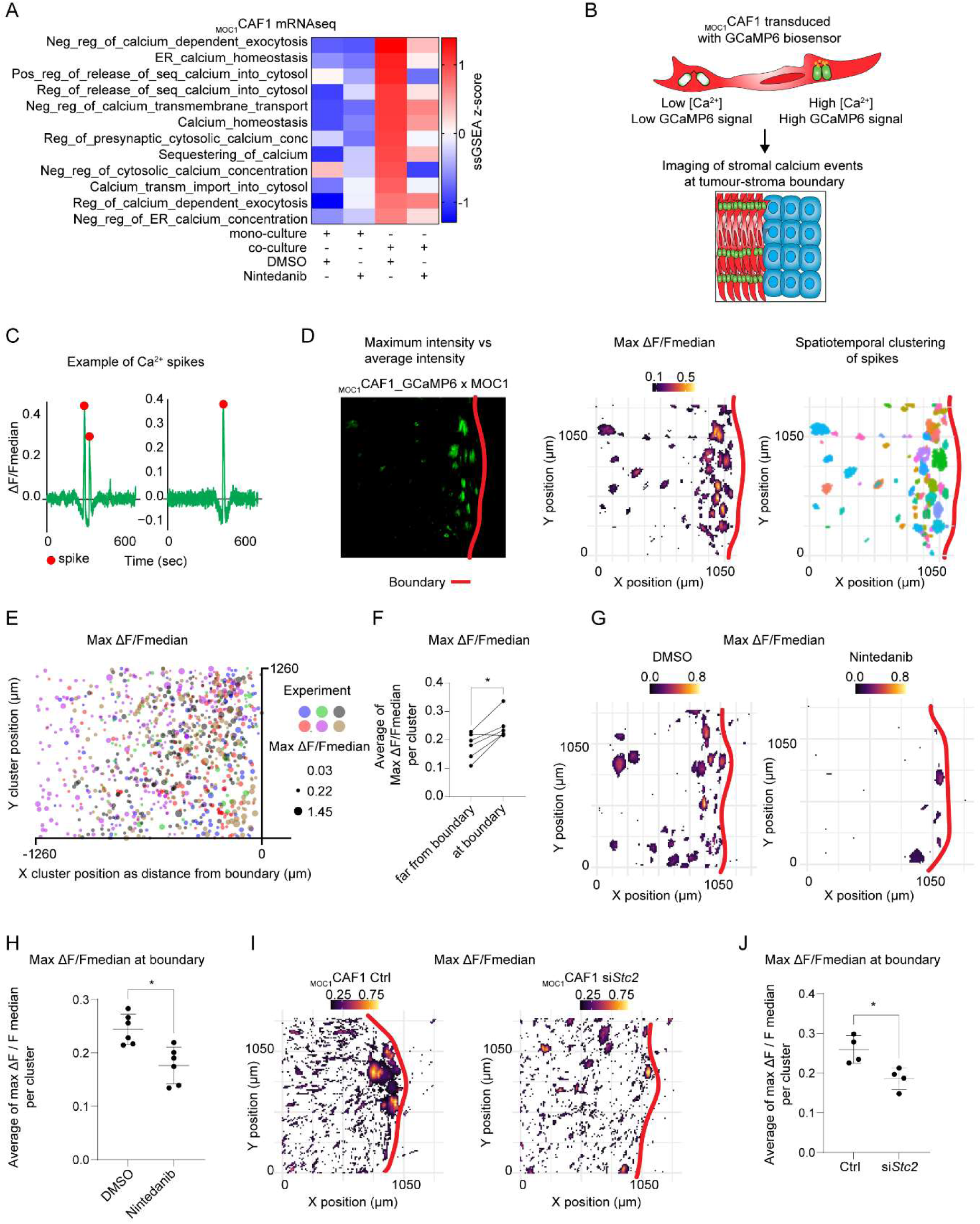
CAF alignment at the boundary boosts multicellular calcium waves. (A) Heatmap of ssGSEA z-score of GO_BP calcium related pathways regulated by nintedanib in co-culture in _MOC1_CAF1. (B) Schematics of GCaMP6 biosensor function and combination with the tumour-stroma boundary assay. (C) Exemplar tracks and spike detection showing ΔF / F_median_ over time. (D) Example of the pipeline used to detect and characterize multicellular calcium spikes at the boundary in time and space. (E) Bubble plot of maximum ΔF / F_median_ spikes plotted with XY coordinated (micron) as a function of distance from the boundary over x axis. Bubble colour represents different biological replicates, bubble size represents maximum ΔF / F_median_ values. (F) Dot plot of average maximum ΔF / F_median_ values far (800-1200 µm) vs at (0-400 µm) the boundary. N=6. Two-tailed ratio paired Student’s t test. (G) XY heatmap of calcium spikes plotting maximum ΔF / F_median_ values for the indicated conditions. (H) Dot plot mean ± SD of average maximum ΔF / F_median_ values at boundary for the indicated conditions. N=6. Two-tailed ratio paired Student’s t test. (I) XY heatmap of calcium spikes plotting maximum ΔF / F_median_ values for the indicated conditions. (J) Dot plot mean ± SD of average maximum ΔF / F_median_ values at boundary for the indicated conditions. N=4. Two-tailed ratio paired Student’s t test.

### Multicellular calcium waves regulate anisotropy and neuronal mimicry in CAFs

We investigated the functional role of intracellular Ca^2+^ in acquisition of CAF anisotropic behaviour and neuronal properties. We used SKF96365 inhibitor, a known drug targeting both store-operated calcium entry (SOCE) and voltage-gated Ca^2+^ channels [42], [43], [44]. Treatment of _MOC1_CAF1 with SKF96365 altered actomyosin cable organization and dramatically reduced alignment (Figure 8A). We confirmed this with 3 human anisotropic cell lines (Figure 8B). In addition, SKF96365 treatment of _MOC1_CAF1 was sufficient to reduce expression of TUBB3 in CAFs and shift the protein to a peri-nuclear sub-cellular localisation (Figure 8C). These data establish the importance of Ca^2+^ release for the higher order organisation of CAFs, and show that two manipulations which reduce coordinated multicellular Ca^2+^ release disrupt CAF alignment at tumour-stroma boundaries. Together, these data predict the existence of CAF-specific MCWs *in vivo* around tumour nests. To test this in a physiological tissue context, we crossed PDGFRA-Cre mice with GCaMP6f reporter mice and performed imaging of live tissue slices with MOC1 late tumours (Figure 8D). This confirmed that CAFs show MCWs around tumour nests (Figure 8E&F and Movie 10&11). In conclusion, we propose that MCWs are fundamental regulator of CAF spatial organization, enabling their higher order anisotropic organisation. This phenomenon is linked to the acquisition of neuronal gene expression programmes, and ultimately leads to CD8+ T cell exclusion and the escape of tumours from immune surveillance (Figure 8G).

**Figure 8:**
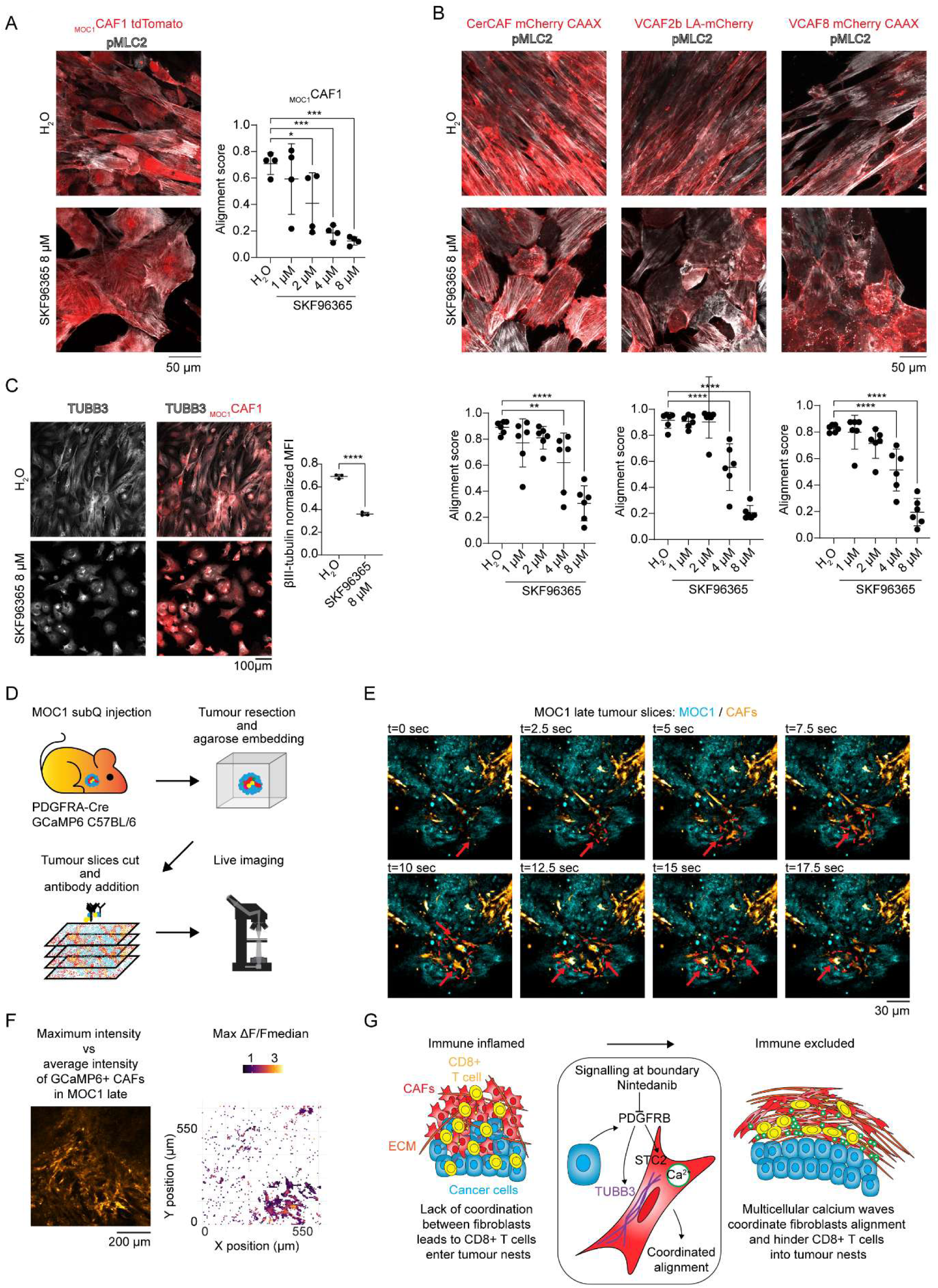
Multicellular calcium waves regulate anisotropy and neuronal properties in CAFs. (A) Left, exemplar images of _MOC1_CAF1 tdTomato and pMLC2 for the indicated conditions. Right, dot plot with mean ± SD of alignment score calculated on tdTomato. N=4. Unpaired one-way ANOVA Holm-Sidak corrected for multiple comparison. (B) Top, exemplar images of human CAF lines and pMLC2 for the indicated conditions. Bottom, dot plot with mean ± SD of alignment score. N=6. Unpaired one-way ANOVA Holm-Sidak corrected for multiple comparison. (C) Left, exemplar images of TUBB3 expression in indicated samples. Right, dot plot with mean ± SD of TUBB3 normalized MFI. N=3. Two-tailed unpaired Student’s t test. (D) Schematics of tumour slices live staining and imaging with a PDGFRA-Cre GCamp6 C57BL/6 mouse. (E) Exemplar time lapse images showing coordinated multicellular calcium burst of GCaMP6-positive CAFs in tumour slices from MOC1 late tumour (MOC1 cells in cyan). Arrows highlight examples of MCW spreading between CAFs. (F) Left, exemplar of maximum intensity over average intensity of calcium spikes in GCaMP6f-positive CAFs in tumour slices from MOC1 late tumour. Right, XY heatmap of calcium spikes plotting maximum ΔF / F_median_ values. (G) Schematics of proposed model.

## Discussion

Tumours are able to evade the immune system via a variety of mechanisms [1]. In this study, we delineate an unexpected functional program that CAFs use to prevent the transit of CD8+ T cells into tumours. As tumours become immune excluded, cancer cell signalling to CAFs increases, with neuronal markers being induced, and increased alignment of the ECM parallel to tumour-stroma boundaries. CD8+ T cells are then guided along matrix fibres parallel to tumour nests, but are unable to access the tumour cells. Through the establishment of a reductionist model of tumour-stroma boundary, we identify a regulatory module controlling the organisation of this barrier to CD8+ T cell access. Nintedanib-dependent and PDGFRB-dependent signals at the boundary promote the expression of a suite of genes, including STC2, that are required for CAF alignment and limiting CD8+ T cell transit. We identify *STC2* as a central player and demonstrate that it regulates MCW at the tumour boundary. Our data suggest that evaluation of the FDA approved drug nintedanib, which is currently used for the treatment of IPF patients [45], in combination with immunotherapy is warranted where stromal barriers prevent CD8+ T cell access to tumours. Indeed, the fact that we observe nintedanib efficacy against tumours rich in well-defined nest structures, but not when mixed tumour-stroma boundaries are present, could enable the identification of patients who would benefit from the drug and explain why other attempts have failed to show a benefit for patients [46], [47]. We propose that the application of metrics of CAF alignment combined with CD8+ T cell staining might be one route to identifying patients likely to benefit from such a combination.

Our observation of MCWs presence indicates that CAFs act in a collective manner. Moreover, this behaviour is enhanced at tumour-stroma boundaries, which is where cancer cell-CAF communication is highest. We propose that MCWs regulate CAF spatial organization, leading to isotropic to anisotropic transition. This helps to ensure optimal coordination of the actomyosin cytoskeleton between CAFs and leads to maintenance of the higher order cell alignment at tumour-stroma boundaries. STC2 is known to regulate Ca^2+^ release from the ER [40]. Actomyosin contractility has been shown to be critical for both the alignment of CAFs and their ability to deposit ECM [25], [48], [49], however no literature – to our knowledge – has shown the importance of MCW in CAF biology. Thus, our work provides a new context and function for Ca^2+^ waves in confluent CAF cultures. Importantly, while virtually all cells change intracellular Ca^2+^ levels, wave-like multicellular propagation of Ca^2+^ are frequently associated with embryonic and neuronal biology [50], [51].

Stromal fibroblasts can adopt a range of sub-states, including ACTA2-positive contractile fibroblasts, IL6-expressing immune-regulatory fibroblasts, and even antigen presenting fibroblasts [2], [3], [19], [24], [52], [53]. Our analyses indicate that the increased alignment and barrier function of CAFs is not linked to any of these states, in line with other reports [25], but is correlated with the up-regulation of the neural marker TUBB3. This links to the growing appreciation of tumours as highly connected entities, most notably in the context of tumour innervation [54], [55], [56]. Recently, acquisition of neuronal properties from cancer cells have been described in lung cancer [57] and melanoma [58]. Our findings clearly demonstrate that neuronal reprogramming is not limited to cancer cells, but can also occur in CAFs. The cues driving this transition are present within the TME and push CAFs into a state of neuronal mimicry. Although fibroblasts and neuronal lineages diverge rather early in development, it is well-established that fibroblasts can differentiate into neurons [59], [60], [61], [62]. Both indirect reprogramming via induced pluripotent stem cell transformation [62] and direct transformation via the use of genetic reprogramming [61] or mechanical stress [59], [60], [61], [62] have been reported. The latter mechanism may be particularly pertinent given the compressive forces present in tumours [63], [64]. We speculate that the expression of neuronal genes enables fibroblasts to align in parallel highly anisotropic arrays similar to neurons [65].

To conclude, our analysis of CAF-mediated immune evasion has uncovered a pharmacologically targetable regulatory signalling network active at tumour-stroma boundaries. We reveal how CAFs undergo isotropic to anisotropic transition via upregulation of MCWs and acquisition of neuronal properties to align parallel to the tumour boundary and limit CD8+ T cell access.

## Acknowledgments

We thank all members of the Sahai laboratory for their comments. We thank the healthy donors’ program and donators for blood donations from the Francis Crick Institute. We thank all the patients participating in the cohort from Bellvitge Biomedical Research Institute in partnership with the Catalan Institute of Oncology. We thank the Cell Sciences, Flow Cytometry, Advanced Light Microscopy, Genomics, Bioinformatics and Biostatistics, Experimental Histopathology, Chemical Biology, Human Biology Facility, Biological Research facility from the Francis Crick Institute for their assistance and training. We thank Lauren Chisholm for support with mouse handling. We thank the Genomics STP, and particularly Deb Jackson, Dan Leonce and Marg Crawford for their contributions to library preparation and sequencing. We thank James Boots for bioinformatic support with bulk mRNAseq analysis. We thank Claudio Ballabio (Leanne Li lab) for GCaMP6s plasmid. We thank Janet Brownlees and Robert Koechl for useful discussions.

## Author contributions

G.G. and E.S. conceptualized and designed the research. G.G. performed the experiments and analysed the results unless otherwise written here below. Z.R. performed the experiments and analysis related to multiplexed IMC and CD8+ T cell migration on top of nanofibres. A.R. performed the experiments related to human A431 / VCAF2b / CD8+ T cell culture and contributed to establish MOC1 MOC2 model in the lab. S.H. supported experimental analysis. D.N. and J.C. provided support with immunostaining of oropharynx HNSCC cohort. H.M. provided support with establishment of 2D tumour-stroma boundary assay. S.L., M.J., M.O. and L.A. provided access to the oropharynx HNSCC cohort. P.C., M.M. D.S. provided bioinformatic support. P.H. and M.A. provided support with IMC. K.H., A.O., X.Y., E.S. provided fundings and access to samples and reagents. G.G. and E.S. wrote the manuscript. All authors reviewed and edited the manuscript. All authors authorized the final version.

## Abbreviations

apCAFs: Antigen presenting CAFs
CAFs: Cancer Associated Fibroblasts
ER: Endoplasmic Reticulum
ECM: Extracellular Matrix
FGFR: Fibroblast Growth Factor Receptor
FN1: Fibronectin 1
FFPE: Formalin Fixed Paraffin Embedded
GOBP: Gene Ontology Biological Processes
IPF: Idiopathic Pulmonary Fibrosis
IMC: Imaging Mass Cytometry
iCAFs: Inflammatory CAFs
IL6: Interleukin 6
HNSCC: Head and Neck Squamous Cell Carcinoma
MFI: Mean Fluorescent Intensity
MAPK: Mitogen-Activated Protein Kinase
MCW: Multicellular Calcium Waves
myoCAFs: Myo-fibroblastic CAFs
NSG: NOD Scid Gamma
PanCK: Pan Cytokeratin
PDGFR: Platelet-Derived Growth Factor Receptor
PDMS: Polydimethylsiloxane
scRNAseq: Single Cell RNA Sequencing
ssGSEA: Single-Sample Gene Set Enrichment Analysis
SD: Standard Deviation
STC2: Stanniocalcin-2
SOCE: Store-Operated Calcium Entry
TGFβ: Transforming Growth Factor-Beta
TME: Tumour Microenvironment
UPR: Unfolded Protein Response
UMAP: Uniform Manifold Approximation and Projection
VEGFR: Vascular-Endothelial Growth Factor Receptor

---

Supplementary Figure 1: Related to Figure 1. (A) Top, R squared of exponential growth is shown. Bottom, average doubling time in days (with 95% confidence interval). N=14 mice. (B) UMAP of scRNAseq entire object split for MOC1 early, MOC1 late and MOC2. N=3 mice per group. (C) UMAP of lymphocyte object. N=3 mice per group. (D) UMAP of stromal object. N=3 mice per group. (E) Box plot with mean ± SD and dots of stromal sub-population proportions over total separated in MOC1 early, MOC1 late and MOC2. N=3 mice per group. (F) Heatmap of probability of communication score for ligands involved in MOC1 late cancer cells – CAF crosstalk via CellChat. (G) Heatmap of gene expression of neuronal-associated genes in CAF dataset.

Supplementary Figure 2: Related to Figure 2. (A) Graph with the 33-marker antibody panel designed and optimised for IMC. (B) Schematics of the 12 major cell types identified with the IMC panel. (C) Left, 10-cell neighbourhoods were defined using the SPATIAL-PHLEX module of TRACERx-PHLEX [33] to identify the nine nearest neighbours of every cell object (‘seed’ cell). Centre, neighbourhood radius plotted as distance to furthest neighbour, average radius of 19.9 µm indicated in red. Right, within-cluster sum of squares plot testing K = 1-10 for 10 cell neighbourhoods to identify optimal number of clusters K. (D) Exemplar images of K means unsupervised clustering of 10 cell-neighbourhoods into K = 4 clusters. Clustering was performed on frequency of cell types in neighbourhoods independent of spatial location and cluster identity was then mapped back onto cell coordinates to plot spatial organisation of cellular communities. (E) Tissue segmentation was performed using a Qupath pixel classifier to segment tissue into tumour and stroma regions based on pixel intensity of epithelial and stromal markers. (F) Heatmap showing the ratio of each indicated cell type in tumour over stroma regions in MOC tissue.

Supplementary Figure 3: Related to Figure 3. (A) Schematics of MOC1 tissue slices preparation, staining and image acquisition. (B) Exemplar images of MOC1 cancer cells (Epcam positive), CAFs (ACTA2-dsRed positive) and CD8+ T cell in MOC1 early and MOC1 late. (C) Dot plot of CD8 MFI over sum of Epcam and DsRed MFI in MOC1 early (N=12) and MOC1 late (N=14). Two-tailed unpaired Student’s t test. (D) Dot plot with mean ± SD of CD8+ T cell track metrics separated in MOC1 early (N=6) and MOC1 late (N=10). Two-tailed unpaired Student’s t test. (E) Dot plot of CD8+ T cell track metrics separated by tumour or stroma in MOC1 early (N=6 per group) and MOC1 late (N=10 per group). Two-tailed paired Student’s t test. (F) TWOMBLI metrics for fibre masks from stromal channels, averaged per sample. N = 13, two-tailed unpaired Student’s t test, p values Benjamini-Hochberg adjusted. (G) Distribution of orientations of tumour boundary and stromal fibres. Representative plot of one image from each tissue type.

Supplementary Figure 4: Related to Figure 3. (A) Representative images of FFPE from oropharynx HNSCC HPV negative patients stained for E-Cadherin, Pan-cytokeratin, ACTA2, Collagen-1, CD8, TUBB3. Green arrows show examples of CD8+ T cells being physically excluded from tumour nests and highlight the length of major nuclear axis being parallel to CAF alignment. (B) Schematics of CAF isolation from tdTomato positive C57BL/6 mice injected with MOC1 or MOC2. (C) Representative image of _MOC1_CAF and _MOC2_CAF positive for tdTomato and stained with ACTA2 and FN1 antibodies. (D) Dot plot with mean of alignment score. N=5 per group. One-way ANOVA Tukey corrected for multiple comparison. (E) Exemplar image of _MOC1_CAF tdTomato positive stained with Collagen1 and DAPI to show spatial correlation between CAF main axis and Collagen fibres. (F) Heatmap of probability of communication score for ligands involved in MOC1 late CAF – CD8+ T cells crosstalk via CellChat. (G) CD8+ T cell migration metrics on different ECM substrates and topologies. Migration metrics extracted using TrackMate [68]. N = 4. Student’s unpaired Student’s t test, p values Benjamini-Hochberg adjusted.

Supplementary Figure 5: Related to Figure 4: (A) Top, exemplar images. Bottom, normalized x axis pixel intensity of CAFs (red) and cancer cell (cyan) channels. (B) Box plot plotting average CD8+ T cell metrics as function of distance from boundary.

Supplementary Figure 6: Related to Figure 4: (A) Exemplar images of tumour-stroma reconstitution assay with addiction of CD8+ T cells using human VCAF2b cervical CAF cell line, human A431 cervical cancer cell line, human primary CD8+ T cells. (B) Example of x axis distribution of CD8+ T cells, VCAF2b and A431. (C) Top, schematics of the assay. Bottom, circular plot of CD8+ T cell directionality at two different distances from the boundary. (D) Exemplar images and the tumour-stroma reconstituted assay with addition of CD8+ T cells showing their tracks and persistence.

Supplementary Figure 7: Related to Figure 5: (A) Left, exemplar images of _MOC1_CAF1 for the indicated conditions. Right, Dot plot with mean ± SD of CAF alignment score. N=3. Two-tailed unpaired Student’s t test. (B) Dot plot mean ± SD of tdTomato positive _MOC1_CAF1 MFI for the indicated conditions as fold change difference over non treated control. N=7. Paired one-way ANOVA Holm-Sidak corrected for multiple comparison. (C) Left, exemplar images of DMSO and nintedanib treated human CAF lines. Right, dot plot with mean ± SD of CAF alignment score. N=5. Two-tailed unpaired Student’s t test. (D) Growth curve of sub-cutaneous injection of MOC1 in C57BL/6 mice. N=16 mice each group. (E) Growth curve of sub-cutaneous injection in NSG mice. N=14 (MOC1 vehicle) and N=15 (MOC1 nintedanib). (F) Growth curve of sub-cutaneous injection of MOC2 in C57BL/6 mice. N=7 mice each group.

Supplementary Figure 8: Related to Figure 6: (A) Heatmap of ssGSEA z-score of HALLMARKS pathway in MOC1 and _MOC1_CAF1 for the indicated conditions. Pathways have been ranked according to how much they are down-regulated in _MOC1_CAF1 co-culture condition with nintedanib versus DMSO. (B) Heatmap of gene expression of top5 UPR genes regulated in _MOC1_CAF1 by nintedanib.

Supplementary Figure 9: Related to Figure 6: (A) Schematics of the strategy used to select CAF-related genes of interest for siRNA screening. (B) Dot plot with mean ± SD of *Stc2* expression in _MOC1_MOCAF1 tested by qRT-PCR. N=3. Two-tailed unpaired Student’s t test. (C) Left, exemplar images of siC- and siSTC2 treated human CAF lines. Right, dot plot with mean ± SD of alignment score of the indicated samples. N=3. Two-tailed unpaired Student’s t test. (D) Kaplan-Meier overall survival (top) and progression free survival (bottom) for HPV negative HNSCC from TCGA datasets stratified for *STC2* expression in high (over median) or *STC2* low (below median). P value was calculated using logRank test. Hazard ratio with 95% confidence interval is shown. (E) Dot plot expression of *PDGFRB* (left) and *ACTA2* (right) log2 mRNA expression in HPV negative HNSCC from TCGA datasets stratified for *STC2* expression in high (over median, N=208 patients) or *STC2* low (below median, N=207 patients). Two-tailed unpaired Student’s t test.

Supplementary Figure 10: Related to Figure 7: (A) Top, dot plot of calcium spikes detected at pixel level showing ΔF / F_median_ values and duration. Bottom, table with indicated metrics. (B) Dot plot of average spike cluster number far (800-1200 µm) vs at (0-400 µm) the boundary. N=6. Two-tailed ratio paired Student’s t test. (C) Bubble plot of number of pixels per cluster plotted with XY coordinated (micron) as a function of distance from the boundary over x axis. Bubble colour represents different biological replicates, bubble size represents number of pixels per cluster. (D) Dot plot of average cluster size (number of pixels per cluster) far (800-1200 µm) vs at (0-400 µm) the boundary. N=6. Two-tailed ratio paired Student’s t test. (E) Bubble plot of cluster duration (seconds) plotted with XY coordinated (micron) as a function of distance from the boundary over x axis. Bubble colour represents different biological replicates, bubble size represents cluster duration (seconds). (F) Dot plot of average cluster duration (seconds) far (800-1200 µm) vs at (0-400 µm) the boundary. N=6. Two-tailed ratio paired Student’s t test.

## Movie legends

Movie 1: exemplar movie of z-projection from a MOC1 early tissue slices from an ACTA2-DsRed C57BL/6 mouse. Slices stained with CD8 antibody (CD8+ T cell) and Epcam (for MOC1 cancer cells). Time frame: 30 seconds.

Movie 2: exemplar movie of z-projection from a MOC1 late tissue slices from an ACTA2-DsRed C57BL/6 mouse. Slices stained with CD8 antibody (CD8+ T cell) and Epcam (for MOC1 cancer cells). Time frame: 30 seconds.

Movie 3: exemplar movie of zoomed-in z-projection from a MOC1 tissue slices from an ACTA2-DsRed C57BL/6 mouse. Slices stained with CD8 antibody (CD8+ T cell) and Epcam (for MOC1 cancer cells). Time frame: 30 seconds.

Movie 4: exemplar movie of zoomed-in z-projection from a MOC1 tissue slices from an ACTA2-DsRed C57BL/6 mouse. Slices stained with CD8 antibody (CD8+ T cell) and Epcam (for MOC1 cancer cells). Time frame: 30 seconds.

Movie 5: exemplar movie of CD8+ T cells migrating when co-cultured with _MOC1_CAF1. CD8+ T cells stained with Far Red Cell Trace dye. Tracks shown with the use of Trackmate [68]. Time frame: 165 seconds.

Movie 6: exemplar movie of CD8+ T cells migrating when cultured with _MOC1_CAF1 and MOC1. CD8+ T cells stained with Far Red Cell Trace dye. Tracks shown with the use of Trackmate [68]. Time frame: 165 seconds.

Movie 7: exemplar movie of CD8+ T cells migrating when cultured with _MOC1_CAF1 and MOC1 treated with DMSO. CD8+ T cells stained with Far Red Cell Trace dye. Tracks shown with the use of Trackmate [68]. Time frame: 165 seconds.

Movie 8: exemplar movie of CD8+ T cells migrating when cultured with _MOC1_CAF1 and MOC1 treated with nintedanib. CD8+ T cells stained with Far Red Cell Trace dye. Tracks shown with the use of Trackmate [68]. Time frame: 165 seconds.

Movie 9: exemplar movie of _MOC1_CAF1-GCaMP6 - MOC1 reconstituted boundary. White line marks boundary. Time frame: 0.2 seconds.

Movie 10: exemplar movie of live tissue slices from MOC1 subcutaneously injected in PDGFRA-Cre GCaMP6 reporter mice. Time frame: 0.2 seconds.

Movie 11: exemplar movie of live tissue slices from MOC1 subcutaneously injected in PDGFRA-Cre GCaMP6 reporter mice. Time frame: 0.2 seconds.

## Methods

### Resource availability

#### Lead contact

Further information should be directed to and will be fulfilled by the lead contact Erik Sahai.

### Materials Availability

Further request for resources, reagents and source data should be directed to and will be fulfilled by the lead contact Erik Sahai.

### Data and Code Availability

#### Data

- scRNAseq have been uploaded on GEO under ID GSE277135 and will be released after acceptance.
- Bulk RNAseq have been uploaded on GEO under ID GSE308232 and will be released after acceptance.

#### Code

This paper did not report original code.

#### Other items

Any additional information required is available from the lead contact upon request.

## Experimental model and study participant details

### Cell lines

MOC1 (EWL001-FP) and MOC2 (EWL002-FP) were purchased from Kerafast. _MOC1_CAF and _MOC2_CAF murine fibroblasts were generated as part of this study. A431 was obtained from the Crick Institute Central Cell Sciences facility. Human fibroblasts (VCAF8, VCAF2B, CerCAF) were already available in the laboratory and obtained as described in [25].

### Mouse models

Table 1 lists all mouse strains used with relevant information.

**Table 1:**
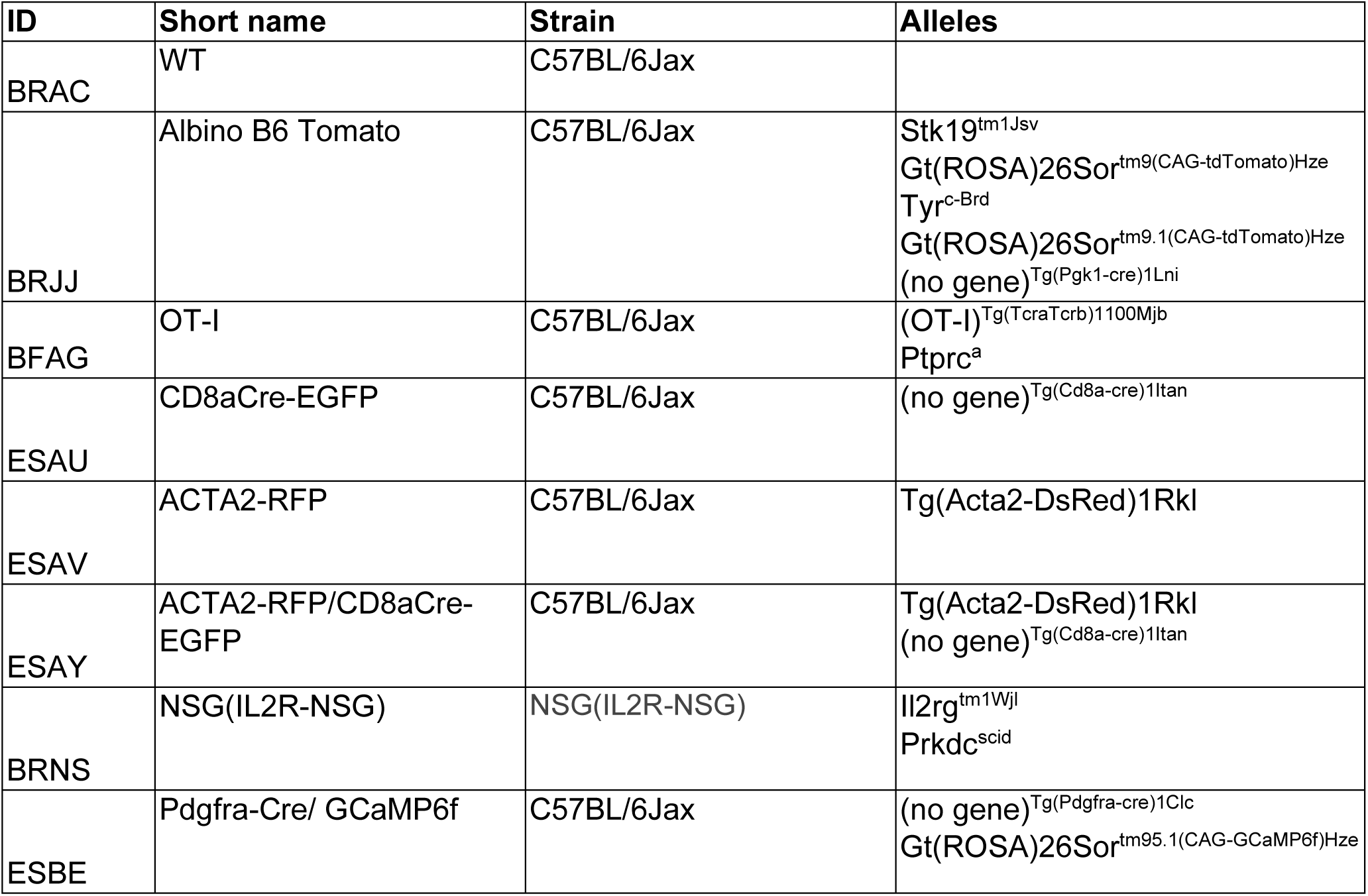
mouse model information.

### Peripheral Blood Mononuclear Cell and monocytes selection

Donations of healthy blood donors were received from the Francis Crick Institute, according to approved protocols of the ethics board of the Francis Crick Institute and the Human Tissue act. Every donor received a Participant Information Sheet and a Consent Form. We do not have access to participant data, including age, sex/gender, ancestry and ethnicity.

Isolated PBMCs were then used to isolate CD8+ T cells.

### TCGA analysis

Analysis performed in Figure S9C&S9D were performed using TCGA web portal ([69]).

### Oropharynx HNSCC cohort

Male and female patients with oropharynx squamous cell carcinoma were enrolled at the Bellvitge Biomedical Research Institute in partnership with the Catalan Institute of Oncology after signature of the fully informed consent. This study was approved by the Research Ethics Committee of Bellvitge University Hospital (Act 04/24), after reviewing all the documentation submitted regarding the research project with Ref. PR138/19. Persona data and confidentiality have been obtained in accordance with Spanish Law 15/1999 on the protection of personal data and Royal Decree 1720/2014 that implements it, as well as Law 14/2007 on biomedical research.

Formalin fixed paraffin embedded (FFPE) samples were collected from patients that underwent curative surgery with or without radiotherapy treatment. Additional information of the patients can be found in Table 2.

**Table 2:**
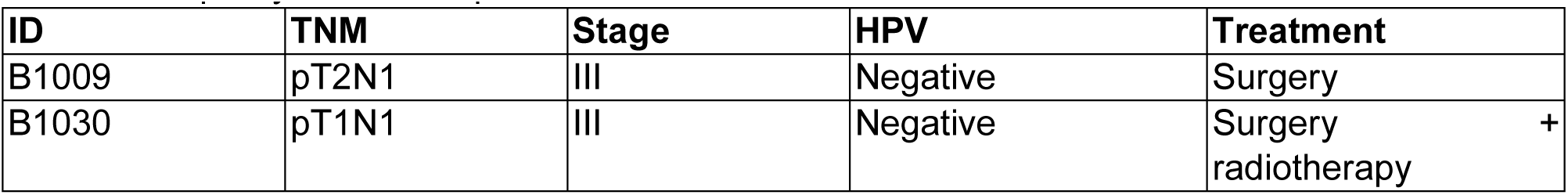
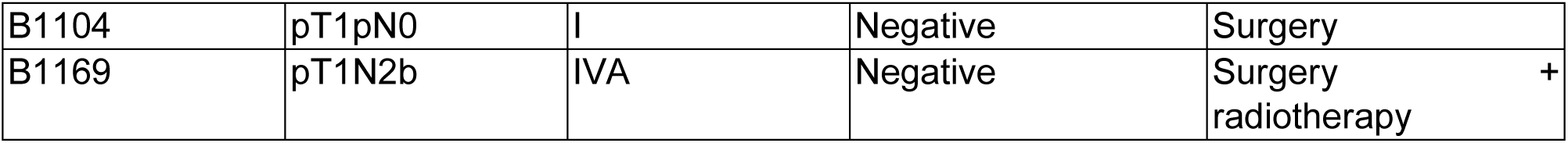
oropharynx HNSCC patient cohort.

| ID | TNM | Stage | HPV | Treatment |
| --- | --- | --- | --- | --- |
| B1009 | pT2N1 | III | Negative | Surgery |
| B1030 | pT1N1 | III | Negative | Surgery<br>radiotherapy |
| B1104 | pT1pN0 | I | Negative | Surgery |
| B1169 | pT1N2b | IVA | Negative | Surgery<br>radiotherapy + |

## Method Details

### Cell lines and reagents

MOC1, MOC2, _MOC1_CAFs and _MOC2_CAFs were cultured in DMEM (ThermoFisher, #41966052) containing 5% fetal bovine serum (Gibco, #10270-106), 1% penicillin/streptomycin (Invitrogen, #15140122), 1% insulin–transferrin–selenium (Invitrogen, #41400045) and kept at 37°C and 5% CO2.

A431, VCAF8, VCAF2B, CerCAF were cultured in DMEM (ThermoFisher, #41966052) containing 10% fetal bovine serum (Gibco, #10270-106), 1% penicillin/streptomycin (Invitrogen, #15140122), 1% insulin–transferrin–selenium (Invitrogen, #41400045) and kept at 37°C and 5% CO2.

Cells were not allowed to reach more than 90% confluency for routine cell culture cultivation. Cell lines that are not commercially obtainable are available from the authors upon reasonable request.

Routine screening for Mycoplasma testing was performed for all cell lines with negative results.

### Cell cultures conditions and treatments

#### Mouse fibroblast isolation and immortalization from MOC tumours

To isolate fibroblasts from MOC1 and MOC2 tumours, we adapted the protocol described by [70]. Briefly, we used a C57BL/6 tdTomato positive mouse and we sub-cutaneous injected MOC1 or MOC2. We dissected tumour masses with a scalpel and placed the tissues into dishes where they were compressed under a 20 mm coverslip pre-treated with HCl 0.1 M over-night. The performed small cuts with the scalpel on the plastic surface of the dishes to increase adhesion of the tissue and favour fibroblast growth. We then covered it with medium DMEM (ThermoFisher, #41966052) containing 10% fetal bovine serum (Gibco, #10270-106), 1% penicillin/streptomycin (Invitrogen, #15140122), 1% insulin–transferrin–selenium (Invitrogen, #41400045). After 5–10 days, fibroblasts started to grow out of the tissue into the coverslip and dish groves. Then, the tissue was removed and the fibroblast population expanded. We discarded wells that were a mixture of fibroblast (tdTomato positive) and cancer cells (tdTomato negative).

For fibroblast immortalization, we used pBABE-HPV E6 vector and selected resistant cells with 2 μg ml^−1^ puromycin.

#### siRNA transfections

RNA interference was performed with Lipofectamine RNAimax reagent from Invitrogen, according to the manufacturer’s instructions. For transient knock down, cells were subjected to reverse transfection with 20 nM RNAi oligos plus forward transfection the day after, then analysed 4-10 days after reverse transfection. Table 3 lists all used RNAi oligos.

**Table 3:** siRNA oligo information.

| siRNA<br>gene<br>name | Species | Catalog# | Gene Accession ID |
| --- | --- | --- | --- |
| ON-TARGET<br>plus Non-<br>targeting<br>Control | Mus Musculus | D-001810-10 |  |
| Aebp1 | Mus Musculus | L-043327-01 | NM_001291857 |
| Angptl1 | Mus Musculus | L-049017-01 | NM_028333 |
| Atp1a1 | Mus Musculus | L-052829-00 | NM_144900 |
| C1qtnf3 | Mus Musculus | L-055527-01 | NM_030888 |
| C4b | Mus Musculus | L-043016-00 | NM_009780 |
| Cdh11 | Mus Musculus | L-053105-00 | NM_009866 |
| Clec3b | Mus Musculus | L-042663-01 | NM_011606 |
| Cnn1 | Mus Musculus | L-060453-01 | NM_009922 |
| Cnn3 | Mus Musculus | L-048093-01 | NM_028044 |
| Col14a1 | Mus Musculus | L-058297-01 | NM_181277 |
| Col16a1 | Mus Musculus | L-057717-01 | NM_028266 |
| Col8a1 | Mus Musculus | L-042577-01 | NM_007739 |
| Cthrc1 | Mus Musculus | L-050227-01 | XM_982785 |
| Cxcl12 | Mus Musculus | L-044397-00 | NM_021704 |
| Cxcl14 | Mus Musculus | L-042905-01 | NM_019568 |
| Dpt | Mus Musculus | L-049523-01 | NM_019759 |
| Ebf2 | Mus Musculus | L-060463-01 | NM_010095 |
| En1 | Mus Musculus | L-051395-00 | NM_010133 |
| Eng | Mus Musculus | L-045109-00 | NM_007932 |
| Fgfr1 | Mus Musculus | L-040832-00 | NM_001079908 |
| Flt1 | Mus Musculus | L-040636-00 | NM_010228 |
| Fndc1 | Mus Musculus | L-053937-01 | XM_985863 |
| Frmd6 | Mus Musculus | L-041225-01 | NM_028127 |
| Gdf10 | Mus Musculus | L-041554-01 | NM_145741 |
| Gem | Mus Musculus | L-047393-01 | NM_010276 |
| Grem1 | Mus Musculus | L-043505-01 | NM_011824 |
| Has1 | Mus Musculus | L-046114-01 | NM_008215 |
| Hhip | Mus Musculus | L-042683-01 | NM_020259 |
| Itga10 | Mus Musculus | L-049662-01 | XM_979750 |
| Itga11 | Mus Musculus | L-041795-01 | NM_176922 |
| Itm2a | Mus Musculus | L-042617-01 | NM_008409 |
| Lgr5 | Mus Musculus | L-045520-00 | NM_010195 |
| Lox | Mus Musculus | L-043942-01 | NM_010728 |
| Loxl2 | Mus Musculus | L-046730-02 | NM_033325 |
| Loxl3 | Mus Musculus | L-059206-01 | NM_013586 |
| Lpar1 | Mus Musculus | L-043497-00 | NM_010336 |
| Lum | Mus Musculus | L-042667-01 | NM_008524 |
| Medag | Mus Musculus | L-040459-01 | NM_027519 |
| Mmp11 | Mus Musculus | L-040421-01 | NM_008606 |
| Olfm2 | Mus Musculus | L-052426-01 | NM_173777 |
| Pappa2 | Mus Musculus | L-044223-01 | XM_355248 |
| Parvb | Mus Musculus | L-042722-01 | NM_133167 |
| Pdgfra | Mus Musculus | L-048730-00 | NM_001083316 |
| Pdgfrb | Mus Musculus | L-048218-00 | NM_001146268 |
| Pi16 | Mus Musculus | L-060227-01 | NM_023734 |
| Piezo2 | Mus Musculus | L-163012-00 | XM_992130 |
| Ptgs1 | Mus Musculus | L-040883-01 | NM_008969 |
| Ptgs2 | Mus Musculus | L-056799-01 | NM_011198 |
| Robo2 | Mus Musculus | L-043024-01 | NM_175549 |
| Rspo3 | Mus Musculus | L-049342-01 | NM_028351 |
| Scara5 | Mus Musculus | L-048174-01 | NM_028903 |
| Sema3a | Mus Musculus | L-058440-01 | NM_009152 |
| Sema3c | Mus Musculus | L-064634-01 | NM_013657 |
| Sema3d | Mus Musculus | L-061000-01 | NM_028882 |
| Serping1 | Mus Musculus | L-043002-01 | NM_009776 |
| Sfrp1 | Mus Musculus | L-048941-01 | NM_013834 |
| Sfrp2 | Mus Musculus | L-042773-01 | NM_009144 |
| Slc16a11 | Mus Musculus | L-054480-01 | NM_153081 |
| Slc26a7 | Mus Musculus | L-053445-01 | NM_145947 |
| Slit2 | Mus Musculus | L-058235-00 | NM_178804 |
| Slit3 | Mus Musculus | L-063651-01 | NM_011412 |
| Spats2l | Mus Musculus | L-059964-01 | NM_144882 |
| Stc2 | Mus Musculus | L-061920-01 | NM_011491 |
| Tek | Mus Musculus | L-045325-00 | NM_013690 |
| Tfpi2 | Mus Musculus | L-062755-01 | NM_009364 |
| Thbs2 | Mus Musculus | L-044025-01 | NM_011581 |
| Tmem204 | Mus Musculus | L-067172-01 | NM_001001183 |
| Tnn | Mus Musculus | L-057883-01 | NM_177839 |
| Tnxb | Mus Musculus | L-044895-01 | NM_031176 |
| Tubb3 | Mus Musculus | L-050529-00 | NM_023279 |
| Wif1 | Mus Musculus | L-046832-00 | NM_011915 |
| Wnt5a | Mus Musculus | L-065584-01 | NM_001256224 |
| Yap1 | Mus Musculus | L-046247-01- | NM_001171147 |

#### Other stable transfections

MOC1 and MOC2 were stably transfected with Lipofectamine 2000 Reagent (Thermo Fisher Scientific) according to the manufacturer’s instructions. Briefly, cell lines were seeded at 2×10^5^ cells in a six-well plate and transfected 1 μg of Piggybac transposase (pPBase-piggyBac) and 1 μg of mTurquoise2 (pPB-mTurq2) plasmid DNAs. After 48h of incubation, the medium with Lipofectamine/plasmid DNA mix was replaced with a fresh medium. Cells were selected using 1 µg ml ^-1^ puromycin. Cells have been further sorted to enrich for population with similar tag expression.

_MOC1_CAF were stably transfected with GCaMP6s in lentiviral vector. Briefly, packaging cells 293T have been transfected with packaging vectors and GCaMP6s plasmid. 48 h later conditioned medium was filtered and applied to _MOC1_CAF cells. Cells have been further sorted to enrich for population with similar tag expression.

Table 4 lists all plasmids used with transfection method details and concentrations used.

**Table 4:** plasmids information.

| Plasmid name | Info |
| --- | --- |
| pPB-mTurq2-T2A-OVA Puro | Available upon request |
| Transposase | Available upon request |
| pGP-CMV-GCaMP6m | Addgene #40754 |

#### Drug treatments

Table 5 lists all drug treatments used with tested concentrations and solvent used.

**Table 5:** drugs information.

| Drug name | Solvent | Concentration | Company | Catalog# |
| --- | --- | --- | --- | --- |
| Afatinib | DMSO | 1µM | Selleckchem | S1011 |
| Dasatinib | DMSO | 0.5µM | LC Labs | d-3307 |
| FAK inhibitor 14 | DMSO | 10µM | Tocris | 3414 |
| GSK269962 | DMSO | 5µM | Cambridge Biosciences | 19180 |
| Lapatinib | DMSO | 1µM | LC Labs | L-4804 |
| Lenvatinib | DMSO | 1µM | Tebu Bio | 282T0520 |
| Nintedanib | DMSO | 1µM | MedChemExpress | HY-50904 |
| Ruxolitinib | DMSO | 1µM | LC Labs | R-6600 |
| SB431542 | DMSO | 1µM | Tocris | 1614 |
| SKF96365 hydrochloride | H<sub>2</sub>O | 8µM | Tocris | 1147 |
| SMIFH2 | DMSO | 10µM | Millipore | 344092 |

**Table 6:**
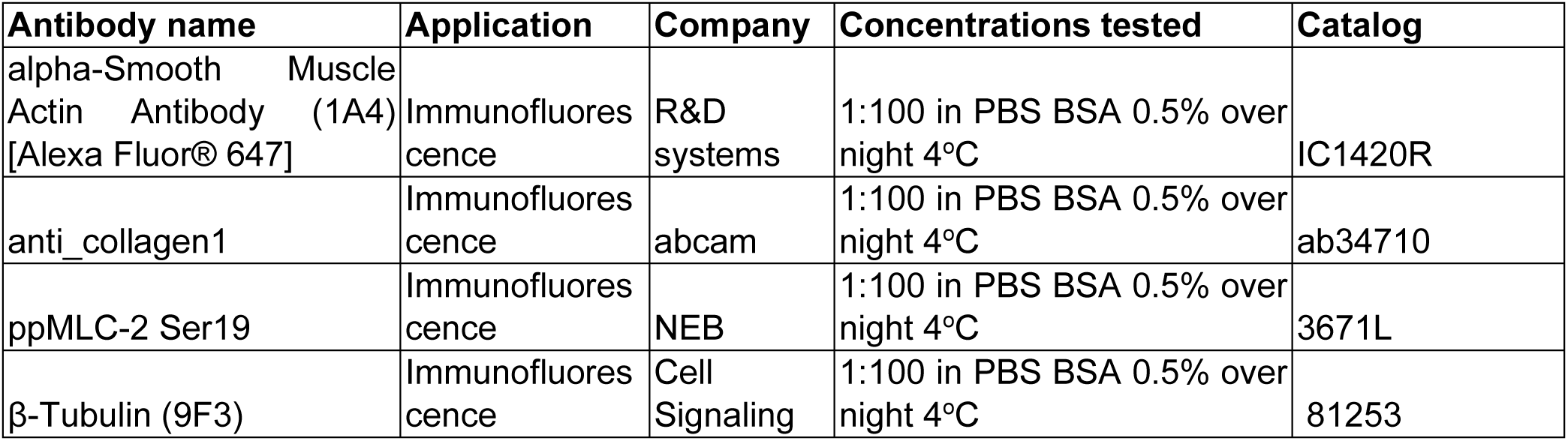
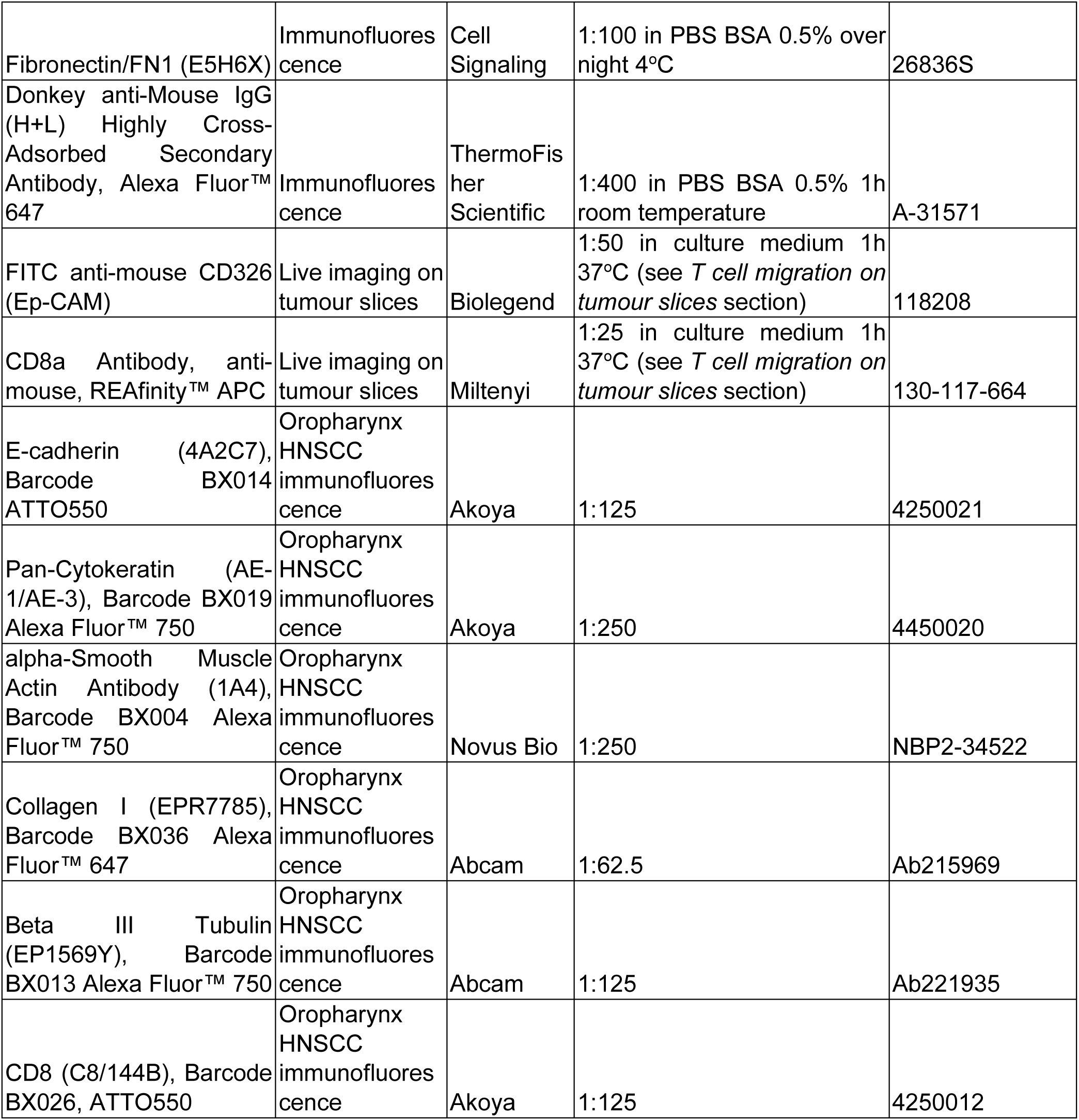
antibodies information.

#### Co-cultures to mimic tumour-stroma boundary

6×10^3^ fibroblasts were seeded in one chamber of a 2-well silicone insert (Ibidi Cat.No:80209) on day 0, in a total volume of 100ul in a 24well. On day 3, medium was changed to fibroblasts and 2×10^4^ cancer cells were seeded in the other chamber. On day 4, the silicone insert was removed and 1 ml medium was added. Depending by assay, co-cultured were kept growing for 2 to 5 days after insert removal.

When T cell dynamics were studied, 5×10^4^ primary CD8 T cells from OT1 splenocytes were added on day 6 using T cell medium.

For calcium imaging, we recorded on day 4/5 after insert removal.

#### Co-cultures and mono-cultures for RNAseq of cancer cells and fibroblasts with or without Nintedanib

Co-cultures were performed with a ratio of 1:2, plating 4×10^5^ fibroblasts and 2×10^5^ cancer cells for a single well of a 6 well plate. For the mono-cultures, we used the same number of cells. Treatment with DMSO or Nintedanib (1 µM, MedChemExpress, # HY-50904) was applied at the time of seeding. Co-cultures and mono-cultures were collected for sorting 24 hours later.

#### Human Peripheral Blood Mononuclear Cell (PBMC) extraction and CD8 T cell selection

Donations of healthy blood donors were received from the Francis Crick Institute. PBMCs were isolated from whole blood using Lymphoprep and SepMate tubes (Stemcell # 07851, and #85450) according to manufacturer’s instructions. Immunomagnetic negative selection EasySep © isolation kits were then used to isolate T cells (Stemcell # 17951) and/or monocytes (Stemcell # 19359). T cells were activated/expanded using CD3/CD28 magnetic beads (ThermoFisher # 11132D at a 1:2 ratio), RPMI media (Gibco #12633 + GlutaMax 1:100 + 10% FBS) containing IL-2 (Peprotech #200-02, final concentration 10 pg ml^-1^) and IL-7 (Peprotech #200-07, final concentration 20pg ml^-1^). Posterior processing depended on the intended experiment.

#### Mouse splenocytes digestion and CD8+ T cell selection

Spleen from OT1 C57BL/6 mice were collected and disaggregated with a 70 µm cell strainer with the help of a plunger from a 2 ml syringe. Plunger and strainer have then been washed to recover all cells with Advanced RPMI with NEAA and Sodium Pyruvate (Thermo Fisher, # 12633012) containing 10% fetal bovine serum (Gibco, #10270-106), 1% penicillin/streptomycin (Invitrogen, #15140122), 1% Glutamax (Life Technologies, #35050061). Splenocytes have then been either frozen or freshly processed to select and activate CD8 T cells. To activate them, 10×10^6^ splenocytes have been incubated with same medium described above supplemented with 100 µM 2-Mercaptoethanol (Thermo Fisher, # 21985023), 5 ng ml^-1^ recombinant murine interleukin-2 (IL2, Peprotech, #212-12-20) and 10 nM SIINFEKL peptide. SIINFEKL peptide was synthetized from Chemical Biology STP at the Francis Crick Institute and resuspended in sterile deionized water. CD8 T cells were used for further experiments between 2-5 days from activation.

### Animal procedures

The Francis Crick Institute’s Animal Welfare and Ethical Review Body and UK Home Office authority provided by Project License 0736231 and 8099471 all animal model procedures. Procedures described in this study were compliant with relevant ethical regulations regarding animal research. For sub-cutaneous injection, MOC1 cells were prepared at 1.5×10^6^ cells per mouse in 100 µl volume made of 70% PBS and 30% matrigel growth factor reduced (Corning, #354230), MOC2 cells were prepared at 0.25×10^6^ cells per mouse with same conditions described for MOC1. For drug studies, Nintedanib (MedChemExpress, # HY-50904) was used as in [71]. Briefly, Nintedanib was resuspended in water with 1% Tween 80 (Sigma-Aldrich) and administered via oral gavage daily at a concentration of 15 mg ml^-1^ for a final dose of 60 mg/kg per mouse. Control animals were dosed with vehicle equivalent to compound dosing groups.

### Oropharynx HNSCC cohort immunostaining and acquisition

Commercially available oligonucleotide-conjugated antibodies were obtained from Akoya Biosciences (Marlborough, MA). In-house oligonucleotide-conjugated antibodies were prepared using the Phenocycler conjugation kit according to the manufacturer’s protocol. All oligonucleotide-conjugated antibodies were stored at 4°C. Complementary fluorophore-conjugated oligonucleotide reporters were also obtained from Akoya Biosciences.

FFPE sections at a 5-micron thickness were mounted onto positively charged slides (Solmedia MSS51012BU). The sections were deparaffinated through two successive 5-minute incubations in xylene. Subsequently, sections were dehydrated through two successive 5-minute incubations in 100% ethanol, followed by a 5-minute incubation in 70% ethanol. Finally, the slides were submerged in distilled water for 1 minute.

Antigen retrieval was performed for 20 minutes using TE buffer (pH 9) in a pressure cooker. After the slides cooled to room temperature, they were rinsed twice in distilled water for 2 minutes each time, incubated twice for 2 minutes in Fusion Hydration Buffer (Akoya Biosciences, 240196), and blocked for 25 minutes in Fusion Staining Buffer (Akoya Biosciences, 240198). The antibody cocktail was prepared in Fusion Staining Buffer supplemented with N-, G-, J-, and S-Blockers (Akoya Biosciences, 7000017), each at 2.375%. The antibody cocktail was then added to the slide under a strip of parafilm and incubated for 3 hours at room temperature.

Following incubation, slides were rinsed three times with PBS, incubated in 1.6% paraformaldehyde (PFA) for 10 minutes, rinsed again three times with PBS, and incubated in ice-cold methanol for 5 minutes. Slides were then rinsed three times in PBS and incubated in the final fixative solution (Akoya Biosciences, 7000017) for 20 minutes. Finally, slides were rinsed three times in PBS and kept at 4°C in Storage Buffer (Akoya Biosciences, 232107) until imaging.

Prior to imaging, the requisite reporter mixes were prepared. Each mix was formulated with 90% 1x Phenocycler buffer (Akoya Biosciences, 7000019), 8.33% Assay Reagent (Akoya Biosciences, 7000002), 1.67% Nuclear Stain (Akoya Biosciences, 7000003), and 5 microliters of each designated reporter. Slides were assembled with a Phenocycler Flow cell (Akoya Biosciences, 240205) and imaged on a PhenoCycler-Fusion system (Akoya Biosciences). The specific antibody concentrations and exposure settings used for imaging are detailed in supplementary Table 6.

### Single cell RNA sequencing (scRNAseq)

#### Tumour dissociation and fluorescence-activated cell sorting for scRNAseq

MOC1 and MOC2 cell lines were injected sub-cutaneous in tdTomato positive C57BL/6 mice. This allowed us to identify cancer cells from host cells.

Tumours were collected and kept in cold DMEM (ThermoFisher, #41966052). Then we performed mechanical dissociation with scissors to relax tumour. Following that, tumours were digested firstly in DMEM containing 5% fetal bovine serum (FBS, Gibco, #10270-106), 1% penicillin/streptomycin (Invitrogen, #15140122), 1% insulin–transferrin–selenium (Invitrogen, #41400045), 1.7 mg ml^-1^ (Merck KGaA/Roche, #11213857001), 0.5 mg ml^-1^ DNaseI (Sigma, #DN-25) for 15 minutes at 37°C with agitation at 200 revolutions per minute (rpm). After that, sequential pipetting with 10 ml and 5 ml pipette was performed, followed by centrifugation at 450 g for 5 min. Pellet was resuspended in 2 ml of TrypLE (Thermo Fisher, #12604013) and kept for 15 minutes at 37°C with agitation at 200 rpm. Following this step, tumours were passed through a 70 µm strainer and washed with DMEM. Tumour digest were then centrifuged at 450 g for 5 min at 4°C.

Pellet was resuspended in FACS buffer: DPBS (no calcium and magnesium, Thermo Fisher, #14190144) with 2% FBS and 1mM EDTA (Thermo Fisher, #15575020). Cells were then centrifuged at 450 g for 5 min. Cells were resuspended in LIVE/DEAD™ Fixable Near-IR (Thermo Fisher, #L10119) and incubated for 15 min at 4°C. Cells were subsequently washed in FACS buffer and centrifuged again at 450 g for 5 min at 4°C. Subsequently, cell were resuspended in FACS buffer containing 1:100 FcR Blocking Reagent (Miltenyi, #130-092-575) for 10 min on ice, followed by centrifuge at 450 g for 5 min at 4°C. Cells were then resuspended with FITC Anti-Mouse CD45-FITC Clone 30-F11 (1:300 in FACS buffer, eBioscience, #103108) for 30 min. Cells were then centrifuged at 450 g for 5 min at 4°C, washed with FACS buffer, re-centrifuged and resuspended in FACS buffer for sorting. For each tumour, we sorted 3 populations: i) tdTomato negative (cancer cells), ii) tdTomato positive and FITC CD45 negative (host cells non immune), iii) tdTomato positive and FITC CD45 positive (host cells immune). Sorted cells were stored in PBS with 2% FBS. We then mixed cells for each tumour at a ratio 1:1:1 to have similar proportions of each category.

Then we centrifuged cells at 450 g for 5 min at 4°C and resuspended in PBS with 0.4% bovine albumin serum (BSA, Acros organics, #240405000) with low binding tubes (Thermo scientific, #90410) at 1000 cells µl^-1^.

#### Cell lyses and library construction protocol

The concentration and viability of the single cell suspension was measured using acridine orange (AO) and propidium iodide (PI) and the Luna-FX7 Automatic Cell Counter. Approximately (3) [5000–20,000] cells were loaded on Chromium Chip and partitioned in nanolitre scale droplets using the Chromium Controller and Chromium Next GEM Single Cell Reagents (CG000315 Chromium Single Cell 3’ Reagent Kits User Guide (v3.1 - Dual Index). Within each droplet the cells were lysed, and the RNA was reverse transcribed. All of the resulting cDNA within a droplet shared the same cell barcode. Illumina compatible libraries were generated from the cDNA using Chromium Next GEM Single Cell library reagents in accordance with the manufacturer’s instructions (10x Genomics, CG000315 Chromium Single Cell 3’ Reagent Kits User Guide (v3.1 - Dual Index)). Final libraries are QC’d using the Agilent TapeStation and sequenced using the Illumina (5) NovaSeq 6000. Sequencing read configuration: 28-10-10-90. Paired-end sequencing.

#### Data preprocessing and quality control

Raw sequencing data was processed into unique molecular identifier count (UMI) matrices using CellRanger (v6.1.2) and a custom built genome incorporating tdTomato sequence (https://www.snapgene.com/plasmids/fluorescent_protein_genes_and_plasmids/tdTomato) to mm10 (GRCm38, version 93), constructed using “cellranger mkref”. Processing beyond this point was carried out in Ubuntu 22.04.3 using R 4.3.1 and the Seurat package (v4.3) [72]. Cells were initially filtered out if: 1) the number of expressed genes was lower than 600, 2) molecular count was lower than 600, 3) more than 20% of UMIs mapped to mitochondrial genes, or 4) they were identified as doublets, using the scrublet package in Python [73]. Only genes that were expressed in at least two cells were retained. After that, the data were normalized, and the top 3000 highly variable genes were detected by the “FindVariableFeatures” function. Next, PCA decomposition was performed and, using the R package “intrinsicDimension” (v1.2) [74], a lower “intrinsic” dimension was estimated for each dataset. The PC within these dimensions were used to construct the UMAP plots for each sample. Further cell exclusion was performed on clusters with high mean mitochondrial content (> 20% of mapped reads). All nine samples in this study were integrated using 3000 variable genes and Seurat’s RPCA method. Clusters were identified for each individual sample and for the integrated samples using “FindClusters” using a range of resolutions [0.1,0.3,0.5,0.7,0.9,1.1] and the Leiden algorithm. In downstream analysis on the integrated samples, only resolutions 0.3 and 0.9 were retained. Cluster-specific gene markers were identified using a Wilcoxon rank sum test, using the function “FindClusters”, and the top 10 genes ranked by logFC per cluster were used to generate a heatmap.

#### Cell-type annotations

Clusters were annotated using the package “Clustifyr” (v1.12) [74] and “scCATCH” (v3.2.2). Clustifyr follows a correlation-based approach on reference transcriptome-signature databases to find an identity with highest similarity for each cluster. Reference databases were pulled from the package “ExperimentHub” (v2.8.1), see Table 7. ScCATCH was used to query our dataset using reference datasets from mouse in tissues “Dermis”, “Epidermis”, “Neonatal skin”, “Skin”, “Submandibular gland”, “Taste bud” and cancer type “Cutaneous Squamous Cell Carcinoma: Skin” [75]. Clusters were further annotated using expression of a priori known markers of interest.

**Table 7:**
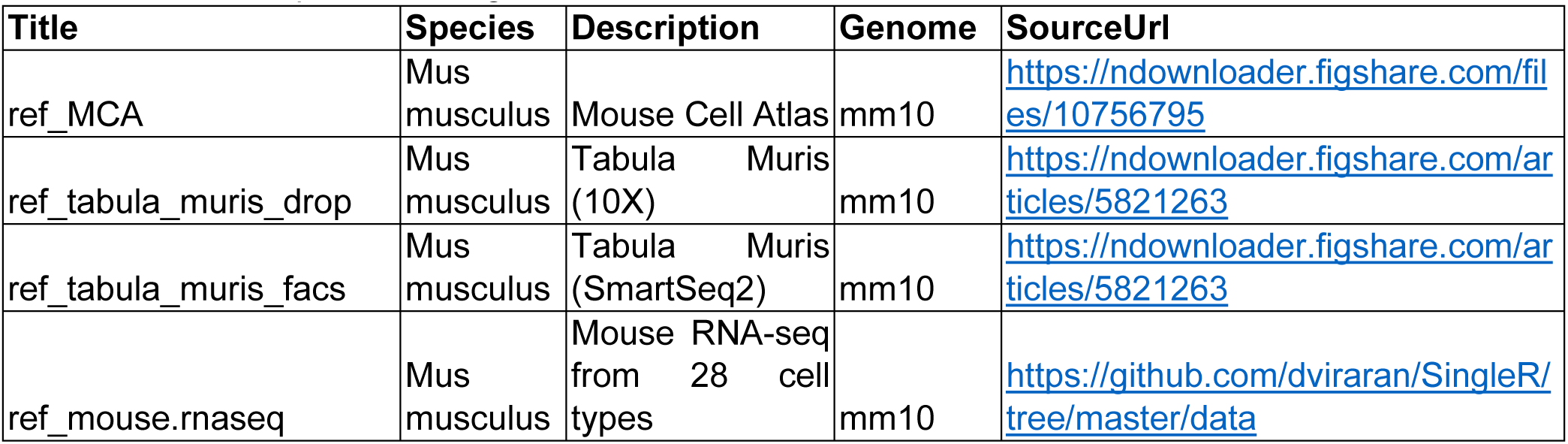
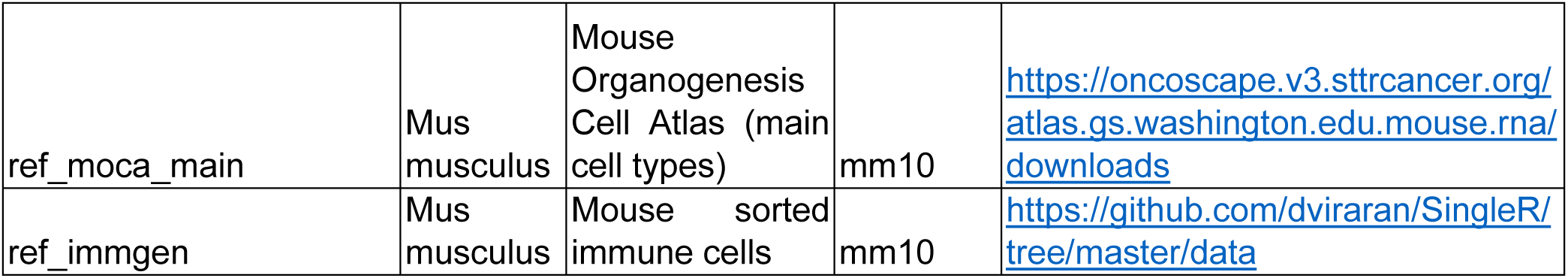
scRNAseq references genomes information.

Further cell-type annotation was performed on the Stromal, Myeloid and Lymphoid cells. After annotation in the integrated dataset, Stromal, Myeloid and Lymphoid cells were individually subsetted, and re-processed using the protocol described in “ScRNA-seq data preprocessing and QC”: scaling, normalisation, top 3000 highly variable genes, PCA, a new UMAP projection, clustering and lead-marker genes per cluster were re-calculated for each of the subsetted datasets. Specific cell-types in the Stromal, Myeloid and Lymphoid cells sub-datasets were identified using the new UMAPs, clustering and lead marker genes per cluster, as well as module scores compiled from custom genes lists using the function “AddModuleScore” from the package Seurat (v4.3).

#### Differential gene expression analysis

Differential gene expression of genes comparing MOC statuses to each other (MOC1E vs MOC1L+MOC2, MOC1L vs MOC1E+MOC2, MOC2 vs MOC1E+MOC1L) was done using GlmGamPoi (version 1.18.0) [76] for selected cell-type of interest in pseudo-bulk manner. To that end, expression profiles were aggregated into pseudobulk expression profiles for each cell type of interest and MOC status group. GlmGamPoi works by implementing a generalised log-linear regression model against a negative-binomial distribution to model the effects of MOC status on cell counts. Results were filtered for 10 > log2 fold change > 0.25 (absolute value) and adjusted p value (Bonferroni correction) < 0.05.

GSEA was carried out on the results for each comparison and cell-type, ranking genes by log2 fold change, using the R package “ClusterProfiler” (v4.8.1) [77], against Gene Ontology (org.Mm.eg.db dataset, https://www.geneontology.org/) and HALLMARKS (https://www.gsea-msigdb.org/gsea/msigdb). Identified gene sets were filtered for pval. < 0.1. To evaluate the expression patterns of genes associated with specific pathways of interest, the VlnPlot function from the Seurat R package was used on either lymphocytes or fibroblast subpopulations. For fibroblast-related analysis, gene sets were retrieved from the Molecular Signatures Database (MSigDB) using the msigdbr R package. Gene Ontology Biological Process (GO:BP) gene sets (category = “C5”, subcategory = “BP”) specific to Mus musculus were extracted and filtered for “calcium” or “neuro” keywords. For each gene set of interest, Seurat’s AddModuleScore function was used to calculate a per-cell aggregated expression score that was visualized using VlnPlot. Statistical comparisons between groups were performed using the Wilcoxon test and annotated on the plots. For CD8 exhaustion marker, the same procedure was used to plot the expression of genes identified as T exhaustion markers in Puram et al [66].

#### Cell–cell communication

Cell–cell interactions based on the expression of known ligand–receptor pairs in different cell types were inferred using CellChatDB (v.1.6.1) [31]. CellChat was run on 1) the whole integrated dataset, between the main annotated cell-types (Cancer, Endothelial, Myeliod, Epithelial, Lymphoid cells, Stromal, RBCs, Melanocytes and Pericytes), and on 2) the Cancer cells, immune and stromal cells sub-types (myoCAFs, iCAFs, apCAFs, Wnt/HH CAFs, iCAFs / myoCAFs, Ki67 iCAFs / myoCAFs, CD8 T cells, Ki67 T cells, CD4 T regs, CD4 T cells, NK cells, Naive / Memory T cells, DN T cells and G/D T Cells). The CellChat analysis was performed as outlined in the CellChat vignette, with the ’population.size’ parameter set to TRUE when computing the communication probability between clusters. Normalized counts from the seurat objects recouping the cell-types of interest were loaded into CellChat, and pre-processed with ‘identifyOverExpressedGenes’, ‘identifyOverExpressedInteractions’ and ‘projectData’ using standard parameters set. The standard ligand-receptor interaction database “CellChatDB.mouse” was selected. For the main analyses the core functions ‘computeCommunProb’, ‘computeCommunProbPathway’ and ‘aggregateNet’ were applied using standard parameters and fixed randomization seeds. Propable communcation between groups of cells were filtered to those detected between cells groups of at least 10 cells (‘filterCommunication’ with ‘min.cells=10’). Finally, to determine the senders and receivers in the network the function ‘netAnalysis_signalingRole’ was applied on the ‘netP’ data slot. Significant interactions were filtered on inter-cluster and intra-cluster interactions were removed.

For comparative analysis, the full dataset was subsetted based on the cell type/line (MOC1 early; MOC1late; MOC2). CellChat was used to determine if there were any differences in ligand-receptor interactions between cell clusters by comparing each cell type/line, with the others [for example MOC1early vs MOC1late + MOC2]. A paired Wilcoxin test was used to determine differential Pathways and ligand-receptor pairs between each cell type relative to non-cell type of interest.

#### STRING analysis on cell–cell communication

To investigate which pathways were involved in cancer cells and fibroblast crosstalk in MOC1 late condition, the cell-cell communication analysis performed on the sub-setted samples was used. Fibroblasts data from myoCAFs, iCAFs, apCAFs, Wnt/HH CAFs, iCAFs / myoCAFs, Ki67 iCAFs / myoCAFs, have been pooled to derive the fibroblast population crosstalk. For each pathway, the sum of all probability of interactions was obtained both for i) “cancer cells communicating to fibroblasts” and for ii) “fibroblast communicating to cancer cells”. For String [67] analysis, the pathways present both in i) and ii) have been used for investigation.

### Imaging Mass Cytometry (IMC)

#### Sample preparation and tumour fixation

MOC1 and MOC2 cell lines were injected sub-cutaneous in tdTomato positive C57BL/6 mice. Half of tumours were digested and processed for scRNAseq and half fixed. Fixation occurred in Formalin solution and samples were embedded into paraffin wax (NBF, Merck KGaA, #HT501128). ROIs for IMC staining were selected from H&E images and 2mm cores of tissue were assembled into a paraffin donor block. Tissue cores were annealed into the donor block with tempering overnight at 43’C followed by 3 cycles of heating to 60’C for 10 minutes and cooling to room temperature for 10 minutes, before heating to 70’C for 10 minutes and leaving to cool completely on a cold plate.

#### Panel design and optimisation

Custom antibodies for IMC imaging were validated against expected staining patterns by standard immunofluorescence staining. Antibodies that gave good signal of expected staining patterns were selected for use in the IMC panel. Custom conjugation to assigned metals was performed using Maxpar-X8 antibody labelling kits (Standard Biotools) according to manufacturer’s protocol. Please see Table 8 for custom antibody catalog numbers and corresponding metal tags.

**Table 8:** heavy metal-tagged catalog and custom antibodies used in IMC staining information.

| Metal | Target | Clone | Company | Catalog # | Staining<br>Dilution |
| --- | --- | --- | --- | --- | --- |
| 141 Pr | aSMA | 1A4 | 3141017D | Standard<br>Biotools | 800 |
| 142 Nd | GFP | Poly | A11122 | Invitrogen | 100 |
| 143 Nd | Vimentin | D21H3 | 3143027D | Standard<br>Biotools | 200 |
| 144 Nd | CD31 | EPR17259 | Ab225883 | Abcam | 200 |
| 145 Nd | CD8 | D4W2Z | 60168SF | CST | 100 |
| 146 Nd |  |  |  |  |  |
| 147 Sm | Tenascin C | EPR4219 | Ab215369 | Abcam | 50 |
| 148 Nd | Pan-Keratin | C11 | 3148020D | Standard<br>Biotools | 100 |
| 149 Sm | CD11b | EPR1344 | 3149028D | Standard<br>Biotools | 400 |
| 150 Nd | TUBB3 | EP1569Y | Ab221935 | Abcam | 200 |
| 151 Eu | Fibronectin | EPR19241-46 | Ab224689 | Abcam | 100 |
| 152 Sm | Pan-CK | AE-1/AE-3 | NBP2-33200 | NovusBio | 200 |
| 153 Eu | F4/80 | D2S9R | 91H030153 | Standard<br>Biotools | 100 |
| 154 Sm | TIM3 | D5D5R | 3154024D | Standard<br>Biotools | 200 |
| 155 Gd | Endomucin | V.7C7 | 14-5851-82 | Thermo | 200 |
| 156 Gd | CD103 | Poly | AF1990 | R&D | 400 |
| 157 Gd |  |  |  |  |  |
| 158 Gd | E-cadherin | 24E10 | 3158029D | Standard<br>Biotools | 100 |
| 159 Tb | Ly6G | E6Z1T | 75028 | CST | 100 |
| 160 Gd | B220 | RA3-6B2 | 14-0452-82 | Ebioscience | 400 |
| 161 Dy | PDGFRa | EPR22059-270 | Ab234965 | Abcam | 100 |
| 162 Dy | CD4 | BLR167J | 91H031162 | Standard<br>Biotools | 100 |
| 163 Dy | FoxP3 | D608R | 72338 | CST | 200 |
| 164 Dy | p-PRAS40 | C77D7 | C77D7 | CST | 200 |
| 165 Ho | Nkp46 | EPR23097-35 | Ab233558 | Abcam | 200 |
| 166 Er | PD-1 | EPR20665 | Ab228857 | Abcam | 100 |
| 167 Er | Podoplanin | 8.1.1 | 14-5381-82 | Thermo | 800 |
| 168 Er | Ki67 | B56 | 3168022D | Standard<br>Biotools | 200 |
| 169 Tm | Collagen 1 | Poly | 3169023D | Standard<br>Biotools | 100 |
| 170 Er | CD3 | Poly | 3170019D | Standard Biotools | 200 |
| 171 Yb | pERK | D12.14.4E | 3171021D | Standard Biotools | 100 |
| 172 Yb | Collagen IV | Poly | NB120-6586 | NovusBio | 100 |
| 173 Yb | CD45 | D3F8Q | 98819SF | CST | 200 |
| 174 Yb | Cleaved caspase 3 | 5A1E | 94530 | CST | 100 |
| 175 Lu | CAIX | Poly | 60168SF | CST | 100 |
| 176 Yb | Histone H3 | D1H2 | 3176023D | Standard Biotools | 100 |

**Table 9:**
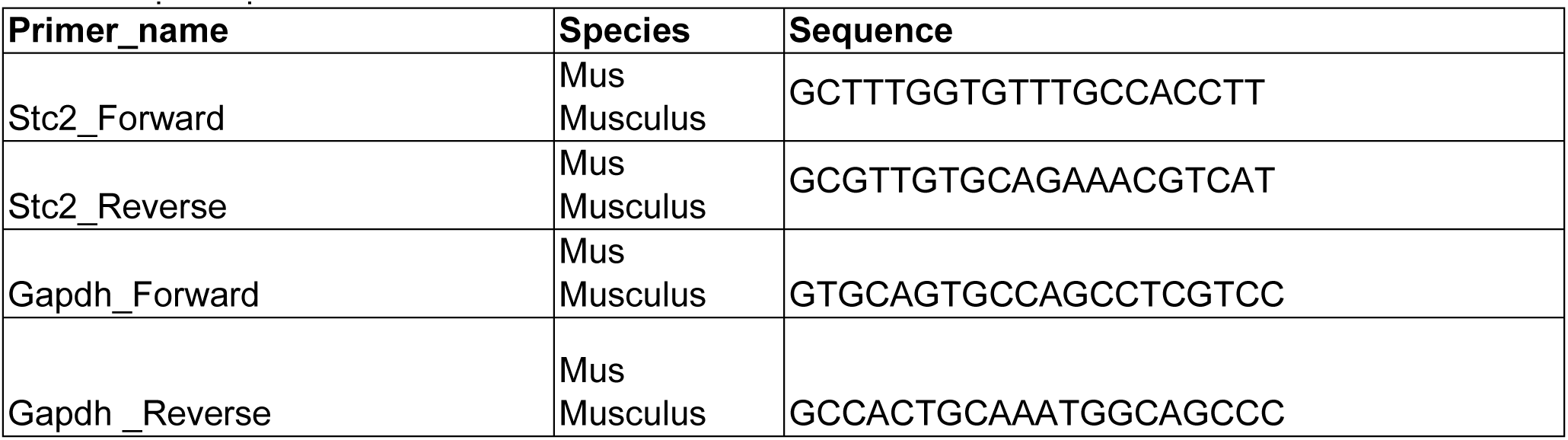
qPCR primers information.

#### Imaging

Imaging was performed as previously described in [78], adapted and optimised for mouse FFPE Tissue. Staining was performed on sections freshly cut 24-48 hours (h) prior to achieve optimum signal. Sections were dewaxed through xylene and microwave antigen retrieval was performed in Tris-EDTA Ph9 buffer (Sigma #T1503, Sigma #E5134) for 22 min at 900W. Sections were blocked for 30 min in Superblock (Thermofisher #35715), and a further 10 min in 1% v/v Mouse Fc-block / PBS-Tween (Miltenyi Biotec, #130-092-575). Slides were incubated with metal-conjugated antibodies (Table 8) in 1% v/v Fc-block / PBS-Tween in a humidified chamber overnight at 4’C. Following antibody staining, slides were stained with Iridium nuclear counterstain (0.2% v/v in PBS, Standard Biotools #2-1192A) for 30 min at room temperature and Ruthenium protein counterstain (0.1% v/v in PBS) for 5 min at room temperature, and left to dry overnight before imaging on the Hyperion CyTOF imaging system.

#### Data analysis

The TRACERx-PHLEX pipeline for highly multiplexed imaging was used for cell segmentation and phenotyping. Cell segmentation was performed using the deep-imcyto ‘Simple Segmentation’ workflow, with nuclear segmentation on Iridium channels and 1-pixel expansion from the nucleus border used to generate cell objects [33]. Cell phenotyping was performed using the TYPEx module, configured according to expression of major lineage markers. Phenotyping quality control was assessed against ground truth cell phenotypes labelled by two scientists. Phenotyping indicated confusion between CD4+ T regulatory cells and CD4+ T helper cells subtypes, so these cells were aggregated to the major type of CD4+ T cells for downstream analysis.

Tissue segmentation was performed using Qupath (v0.4.3). A pixel classifier was trained to classify tissue into tumour, stroma or background regions based on expression of epithelial and stromal marker channels. Segmented objects were exported to Fiji-ImageJ (v2.14.0) for pixel assignment to regions. Cells were matched to classified pixels based on centroid X/Y coordinates in R (RStudio version 4.3.1). Cell density for whole tissue and segmented regions was calculated normalised to total tissue / region area (from Qupath segmented objects).

10-cell neighbourhoods were identified using the Spatial-PHLEX module of TRACERx-PHLEX to calculate the nine nearest neighbours of every cell based on cell centroid coordinates [33]. Neighbourhood interactions were normalised to the density of the seed cell of interest. K-means unsupervised clustering of neighbourhoods based on cell type frequency was performed using the ClusterR package in R (RStudio version 4.3.1).

Further data analysis and plotting was performed in R (RStudio version 4.3.1).

#### TWOMBLI fibre analysis

Fibre analysis of ECM and aSMA channels was performed using the TWOMBLI plugin for ImageJ [34]. Individual IMC channels were de-speckled prior to analysis to remove noise and thresholded to remove edge artifacts. Fibre masks were segmented using a minimum line width of 5, maximum line width of 10, minimum branch length of 10, maximum curvature window of 20, minimum curvature window of 10, curvature window step size of 10, contrast saturation of 0.35 and minimum gap diameter of 5. For comparison of fibre orientation relative to the tumour boundary, boundary masks were generated by expanding tumour and stroma masks from Qupath segmentation by 1 pixel and creating a boundary mask from the overlapping pixels. The boundary masks and TWOMBLI fibre masks were tiled into 200 µm ROIs and the Dominant Direction tool of the Orientation-J ImageJ plugin was used to calculate the major angle of every ROI. Circular correlation between angles was calculated using the ‘circular’ package in R.

#### mRNA sequencing analysis for co-cultures

*Cell sorting and RNA extraction*

After 24 h from cell seeding, _MOC1_CAF and MOC1 cells were treated with Trypsin-EDTA (Thermo Fisher, #25200056) and immediately centrifuged at 300 × g for 5 min to remove supernatant. Cells were then resuspended in FACS buffer: DPBS (no calcium and magnesium, Thermo Fisher, #14190144) with 2% FBS and 1mM EDTA (Thermo Fisher, #15575020). Cells were then centrifuged at 450 g for 5 min.

From each experiment and condition, we sorted 2 populations: i) MOC1, positive for mTurquoise2; ii) _MOC1_CAF, positive for tdTomato.

Sorted cells were then lysed with 350 µl RLT buffer (Qiagen, 79216) containing 1% β-mercaptoethanol (Sigma, M6250). Total RNA was extracted using the RNAeasy Mini kit (Qiagen, 74104; n = 3 independent experiments). Prior to library construction, the quality of total RNA was assessed by Bioanalyzer 2100 (Agilent Technologies Inc).

#### mRNAseq processing

Bulk mRNAseq data was processed using the Nextflow (v23.10.0) nf-core-rnaseq pipeline (v3.10.1-g6e1e448)[79]. Reads were aligned to the Ensembl Mus_musculus.GRCm39.112 genome, using STAR 2.7.10a and quantified using RSEM 1.3.1. Normalisation of raw count data (vst normalisation) was performed with the DESeq2 package [80] (v1.42.1) within the R programming environment (v4.3.2) and was used to plot expression of single genes of interest.

#### Gene set enrichment analysis (GSEA)

GSEA was performed with GSEA software v4.3.3. The dataset used to perform the comparative analysis were murine Hallmarks (mh.all.v2025.1.Mm.symbols.gmt) [81]. All the parameters have been used as defaults except: permutation type (gene set) and metric for ranking genes (Student’s t-test).

#### Single sample GSEA (ssGSEA)

ssGSEA was performed with R package GSVA [82] and searching for gene signatures downloaded MsigDB [81]: murine Hallmarks (mh.all.v2025.1.Mm.symbols.gmt), GO:BP all signatures and GO:BP gene sets related to “calcium” keywords. Subsequently, per sample z-score normalization has been applied. For calcium analysis, gene signatures with fewer than 7 genes have not been further analysed. Figure 7A shows those gene signatures found to be upregulated in _MOC1_CAF co-culture and downregulated by Nintedanib.

### Live imaging

#### T cell migration on tumour slices

MOC1 cells were injected sub-cutaneous in ACTA2-dsRed C57BL/6 mice, which allowed us to observe *ex vivo* ACTA2-positive fibroblasts. Once the tumours have been harvested and stored in cold PBS, have been added into a mold with low melting agarose 4% (Sigma, #A9414), then added immediately the tumour inside. After that, we waited 30 min for agarose to solidify. Following this, mold was removed and the agarose was glued to the flat surface to circular disc, which then was added to the rectangular case and placed into Leica VT1200S, surrounded with ice and filled with cold PBS. Wilkinson blasé was added. Slices were set at 1mm/s speed and 2.5mm amplitude. Slices were cut 350 µm thick.

For antibody staining, slices were incubated for 30 min at 37°C with T cell medium (described in *Mouse splenocytes digestion and CD8 T cell selection*), with addition of 1:100 FcR Blocking Reagent (Miltenyi, #130-092-575). Subsequently, we used the same medium with addiction of CD8a-APC (Miltenyi, #130-117-664) and CD326-FITC (Biolegend, #118208) and incubated slices for 1h at 37°C. After this step, slices have been flattened on the 24-well glass-bottom dishes (MatTek, #P24G-1.0-13-F) and covered with low melting agarose 1%. After solidification, T cell medium was added and live imaging was performed the same day.

For imaging acquisition, slices were acquired with confocal microscope at 37 °C and 5% CO_2_ (LSM 980 equipped with environmental chamber and CO_2_ mixer). Images were takes with ×10 Plan-APOCHROMAT, NA 0.45, Zeiss objective. For time lapses, images have been acquired every 30 sec for at least 30 min.

T cell tracking was performed using TrackMate ImageJ plugin (v7.11.1) [68] with the following parameters: for spot detection LoG despot was used selecting for spots with estimated object diameter 10µm and filtering spots with initial quality below 0.45. For tracks detection, Simple LAP tracker was used with linking max distance of 15 µm, gap-closing max distance of 15 µm and gap-closing max gap frame of 2. Tracks lasting less than 270 sec have not been further analysed.

*T cell migration with co-cultures to mimic tumour-stroma boundary and on monolayers* Activated murine CD8 T cells (from OT1 C57BL/6 mice) or human CD8 T cells (from helathy donors) were dyed using CellTrace Far Red (ThermoFisher Scientific, #C34564) and plated onto tumour-stroma boundaries or monolayers at 5×10^4^ cells per well and left to rest for 5/6 h. For imaging acquisition, samples were acquired with epifluorescence time-lapse imaging at 37 °C and 5% CO_2_ (Nikon Ti2 inverted microscope fitted with an Okolab environmental chamber and CO_2_ mixer). Images were takes with ×10 Plan Fluor, NA 0.3 Ph1, Nikon objective. For time lapses, images have been acquired every 165 sec for at least 5 hours.

T cell tracking was performed using TrackMate ImageJ plugin (v7.11.1) [68] with the following parameters: for spot detection LoG despot was used selecting for spots with estimated object diameter 10µm and filtering spots with initial quality below 0.45. For tracks detection, Simple LAP tracker was used with linking max distance of 50 µm, gap-closing max distance of 15 µm and gap-closing max gap frame of 2. Tracks lasting less than 14850 sec have not been further analysed.

#### T cell migration on nano-fibres

To model T cell migration on ECM fibres, random and aligned nanofibre mats (Sigma Aldrich-Merck KGaA, #Z694517 & #Z694614) were coated with substrate proteins; Collagen-1 (Corning #354249) and Fibronectin (Sigma, #F1141-1MG) at 5 µg/cm2 before blocking with 2% BSA in PBS. Mats were attached to 12 well plates using 2% w/v agarose gel (ThermoFisher Scientific, #16500500).

Activated murine CD8 T cells (from OT1 C57BL/6 mice) were dyed using CellTrace Far Red (ThermoFisher Scientific, #C34564) and plated onto ECM-coated mats at 6×10^4^ cells per well. For imaging acquisition, samples were acquired with epifluorescence time-lapse imaging at 37 °C and 5% CO_2_ (Nikon Ti2 inverted microscope fitted with an Okolab environmental chamber and CO_2_ mixer). Images were takes with ×10 Plan Fluor, NA 0.3 Ph1, Nikon objective. For time lapses, images have been acquired every 120 sec for at least 2 hours.

T cell tracking was performed using TrackMate ImageJ plugin (v7.11.1) [68] with the following parameters: for spot detection LoG despot was used selecting for spots with estimated object diameter 15µm and filtering spots with quality above 1. For tracks detection, Simple LAP tracker was used with linking max distance of 50 µm, gap-closing max distance of 50 µm and gap-closing max gap frame of 2. Tracks lasting less than 6600 sec have not been further analysed.

#### Calcium pulses acquisition and data analysis

For image acquisition of calcium pulses acquisition with tumour-stroma boundaries or monolayers, samples were acquired with epifluorescence time-lapse imaging at 37 °C and 5% CO_2_ (Nikon Ti2 inverted microscope fitted with an Okolab environmental chamber and CO_2_ mixer). Images were takes with ×10 Plan Fluor, NA 0.3 Ph1, Nikon objective. For time lapses, images have been acquired every 0.2 sec for at least 10 min.

For image acquisition of calcium pulses acquisition with MOC1 tumour slices, MOC1 cells were injected sub-cutaneous in PDGFRA-Cre GCaMP6f C57BL/6 mice. Slices were obtained as in “T cell migration on tumour slices” and stained only with CD326 antibody. Slices were acquired with confocal microscope at 37 °C and 5% CO_2_ (LSM 980 equipped with environmental chamber and CO_2_ mixer). Images were takes with ×10 Plan-APOCHROMAT, NA 0.45, Zeiss objective. For time lapses, images have been acquired every 0.2 sec for at least 10 min.

Spikes detection was obtained was R package. Calcium imaging data were acquired as .txt files containing mean fluorescent intensity (MFI) values for each pixel over time. MFI was normalized by dividing each pixel’s intensity by the average intensity of all pixels at the same time frame (global normalization). To correct for baseline drift and smooth local fluctuations, a LOESS (locally estimated scatterplot smoothing) regression was applied to each pixel time series using a smoothing span of 0.3. The normalized signal was divided by the LOESS fit to obtain a detrended and corrected fluorescence trace. For each pixel, the baseline fluorescence was estimated as the median of the corrected signal across time, and the noise level was estimated as the standard deviation of that signal. A spike was defined as any time point at which the signal exceeded a threshold of baseline + 5 × standard deviation.

To avoid detecting multiple points within a single spike as separate events, nearby spike detections at the same pixel were merged if they occurred within 100 frames (2 seconds) of one another. The final spike time was assigned to the frame with the maximum amplitude within each group. For each spike, the ΔF/F_median_ (deltaF_over_F_median_) was computed as: ΔF/F_median_ = (peak amplitude − baseline) / baseline.

The duration of each spike was also calculated as the time (in milliseconds) for which the corrected signal remained above half of its peak amplitude within a ±100-frame window around the spike.

To identify spatiotemporal clusters of activity, spike events from all pixels were analysed using density-based spatial clustering (DBSCAN). Each spike was represented in a 3D coordinate space defined by its spatial position (X, Y) and normalized time (Tn = Time / 10). This normalization makes 10 frames equivalent to 1 unit in time, balancing spatial and temporal proximity. DBSCAN was run with parameters: eps (maximum distance to consider points in the same neighbourhood) at 2 and minPts (minimum number of points required to form a cluster) at 3. For each detected cluster, the number of involved pixels, total duration, maximum ΔF/F_median_, and the cluster’s spatial centre were calculated.

### Alignment

T cell track alignment and fibroblast alignment was measured with Directionality ImageJ plugin for all figures expect for Figure 3I&3J&3K&S3F&S3G, where OrientationJ ImageJ plugin was used.

### Boundary height score

To measure the physical height of the tumour-stroma boundary, fibroblast MFI values over x axis were extracted. Then, MFI values have been organized from as follows: a) extracted the average of top 2% MFI values over x axis, representing the values around the boundary; b) extracted the average of the 2% MFI values further away from the boundary. The height score represents the ratio of a over b.

### T cell boundary accumulation score

To measure T cell boundary accumulation, CD8 T cell MFI values were thresholded and then values over x axis were extracted.

Then, MFI values have been averaged in 10 tiles according to the x position (i.e. first 10% x axis values, second 10% x axis values, etc…). For each sample, these values have been organized from as follows: a) sum of each tile, representing the total amount of T cell MFI values; b) maximum value of this tile, representing the tile with higher number of T cells.

The T cell boundary accumulation score represents the ratio of b over a.

### Immunofluorescence assay

Samples were fixed for 10 min at room temperature with PFA 4%, then washed twice in PBS. Permeabilization was obtained by incubating with 0.5% Triton X-100 in PBS, blocked (3% BSA in PBS) and incubated overnight at 4 °C in primary antibodies diluted in 0.5% BSA in PBS. Cells were then washed before incubating in secondary antibody diluted in 0.5% BSA in PBS for 1 h. Cells were washed before adding DAPI at 1:1,000 (Sigma, D9542) and imaged using a Zeiss LSM980 microscope. Images were analysed using ImageJ.

Table 6 lists all antibodies used with tested concentrations and relevant infos.

### RNA extraction and RT-qPCR

Cells were collected and lysed with RLT buffer and total RNA was extracted using the RNeasy Mini kit (Qiagen, #74104), according to the manufacturer’s protocol.

The cDNA was prepared using M-MLV reverse transcriptase (Promega, #M3682), and quantitative PCR was performed using PowerUp™ SYBR™ Green Master Mix (ThermoFisher, #A25778), using the QuantStudio 3 and 7 Real-Time PCR systems (Applied Biosystems).

Custom primers were acquired from Sigma; sequences are available in Supplementary Table 9. RNA levels were normalized using three house-keeping genes using the ΔΔC method and reported as relative fold change compared with control.

### Software and visualization

Graphs were generated with Prism software (Graphpad Software v10.5.0), R studio (v4.2.1 or v2025.5.1), Adobe Illustrator 2024 (v28.3). scRNAseq data were analysed with Seurat package (v4). FACS data were analysed with FlowJo software (v8). Micrograph data have been handled with ImageJ (v1.54p) or MCD Viewer (v1.0.560.6) for IMC data. Oropharynx HNSCC cohort immunofluorescence, whole slide images were visualised and representative regions-of-interest exported using QuPath (v5.0.0). GSEA analysis has been performed with GSEA software (v4.3.3). References have been organized with Mendeley Reference Manager (version 2.116.0, Elsevier Ltd).

### Quantification and statistical analysis

Statistical analysis was performed using Prism software (Graphpad Software v10.5.0), Excel software (Microsoft Corporation v2108), RStatix package in R studio (v4.2.1 or v2025.5.1). All statistical tests performed are explained in the corresponding figure legend. All N values are biological replicates. P values are represented as: *, p < 0.05, **, p < 0.01, ***, p < 0.001, ****, p < 0.0001.

