## Supplementary figures for "Multicellular Calcium Waves in Cancer-Associated Fibroblasts Regulate Neuronal Mimicry and Anisotropy Leading to Immune Exclusion"

Supplementary Figure 1

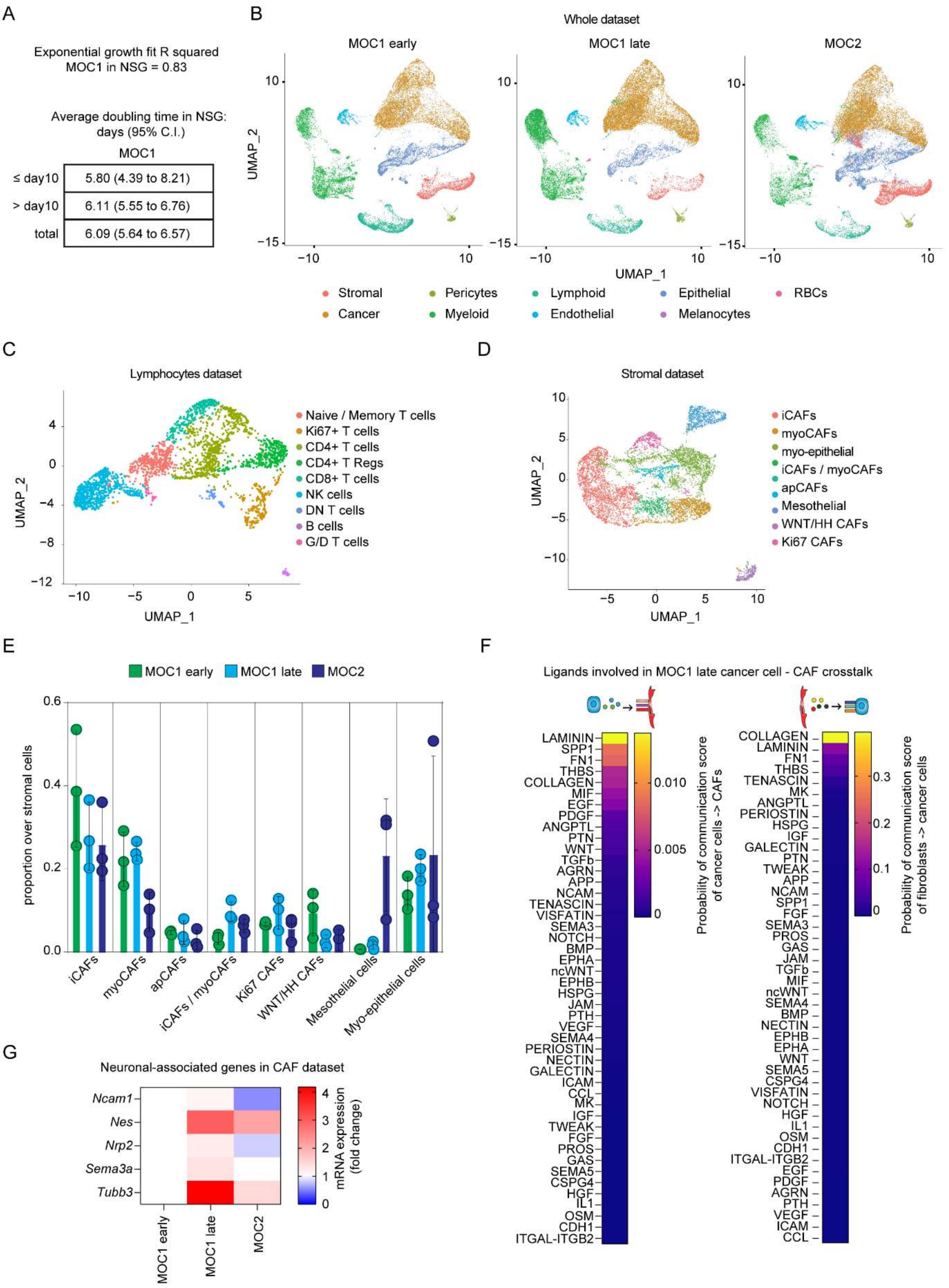

Supplementary Figure 2

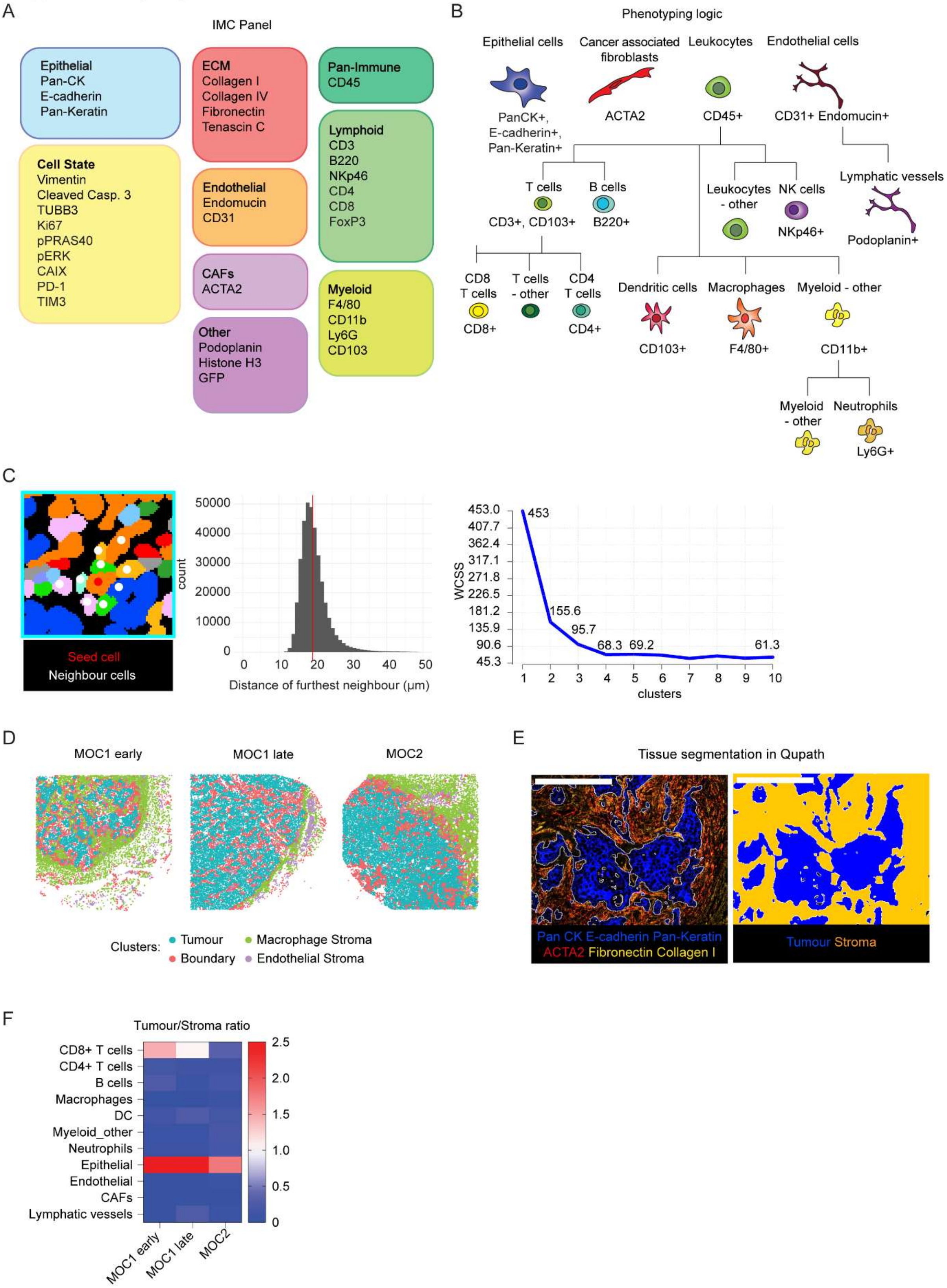

#### Supplementary Figure 3

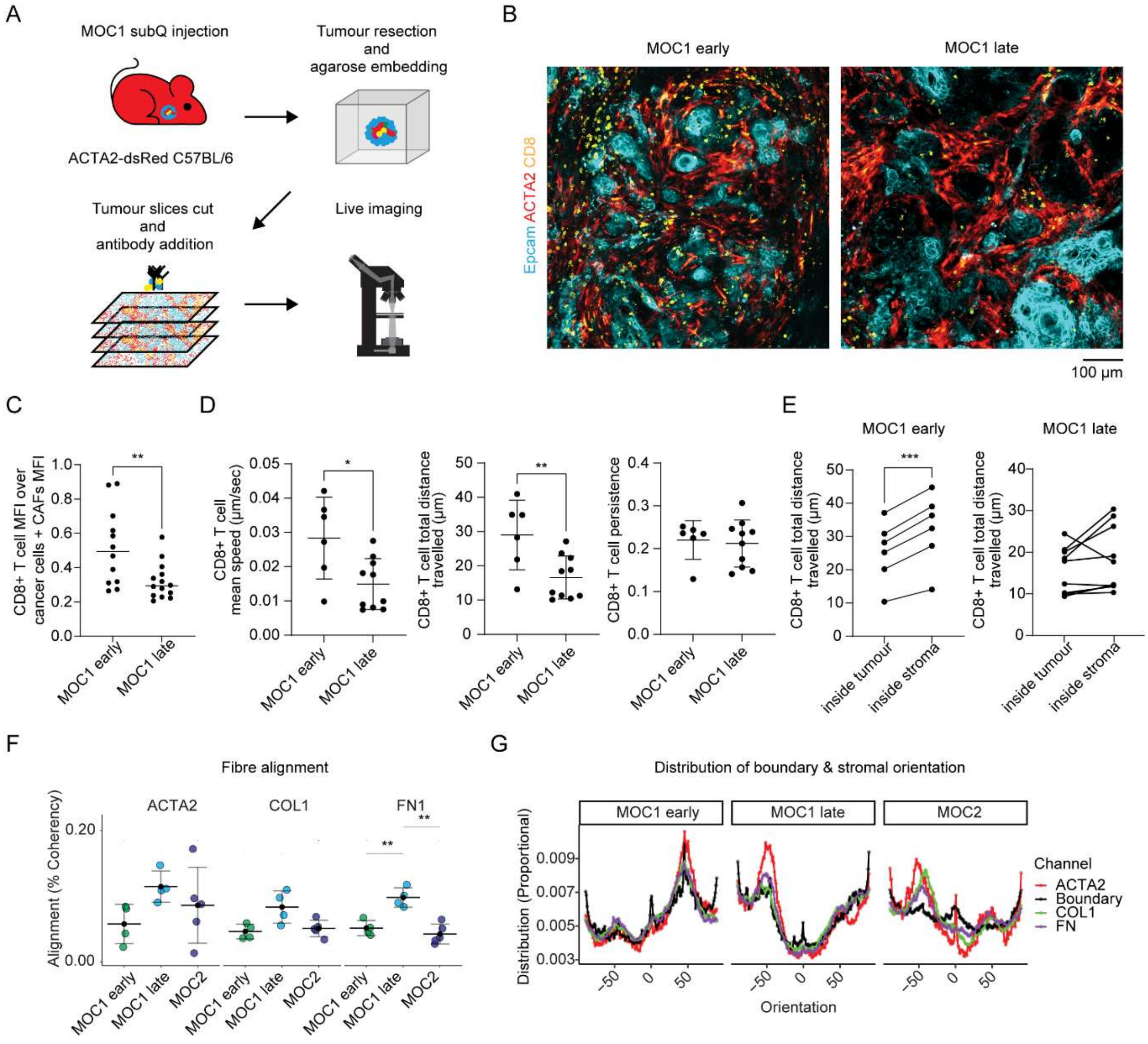

Supplementary Figure 4

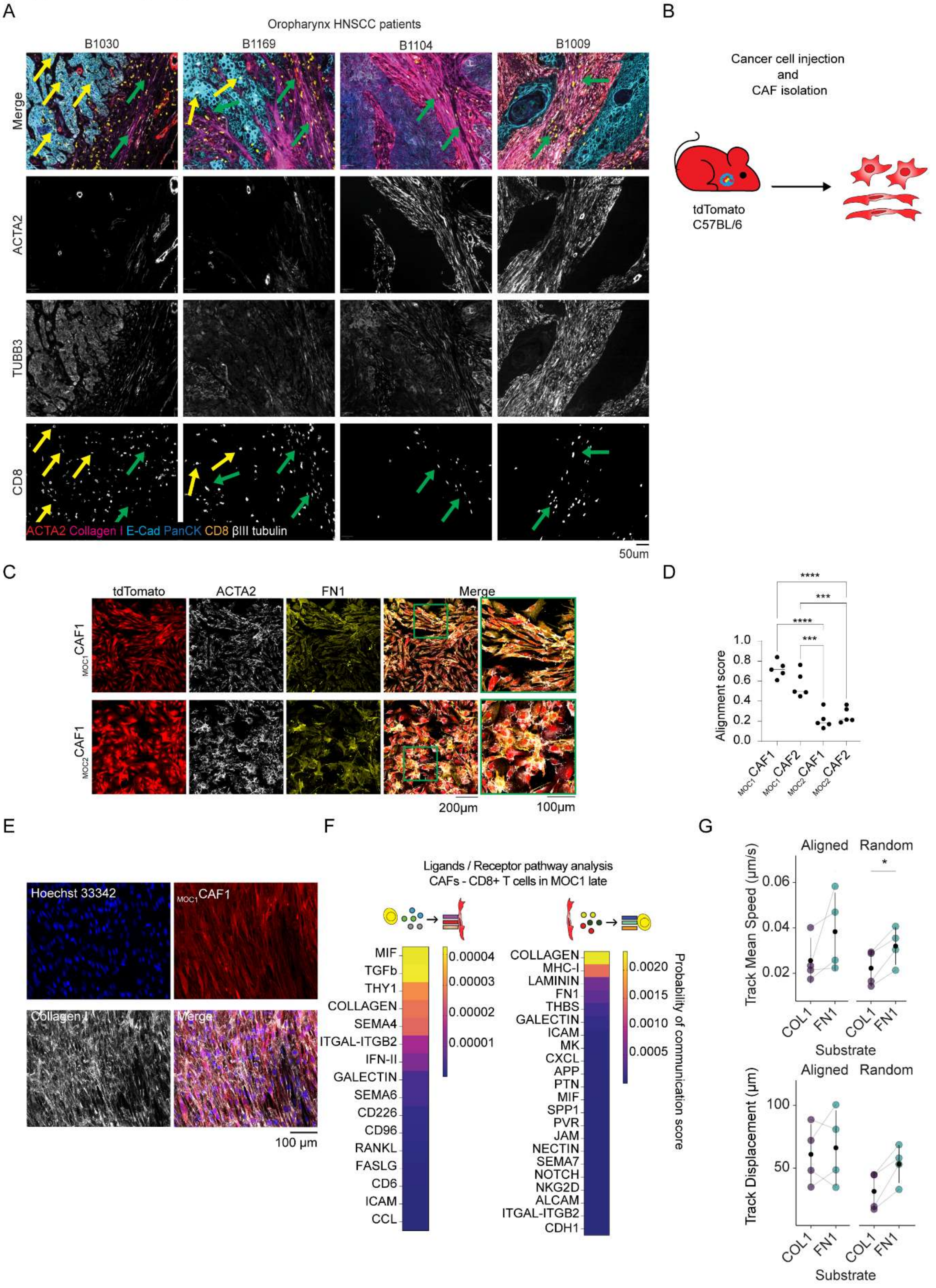

### Supplementary Figure 5

A

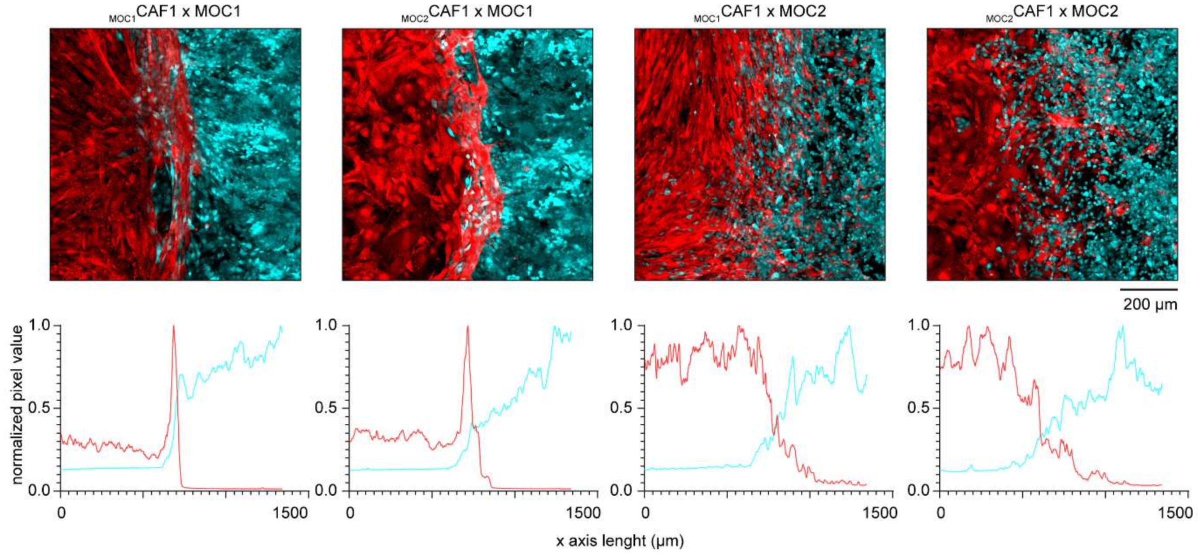


B

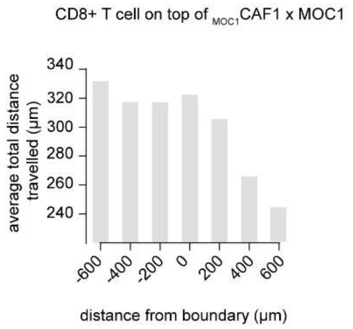

Supplementary Figure 6

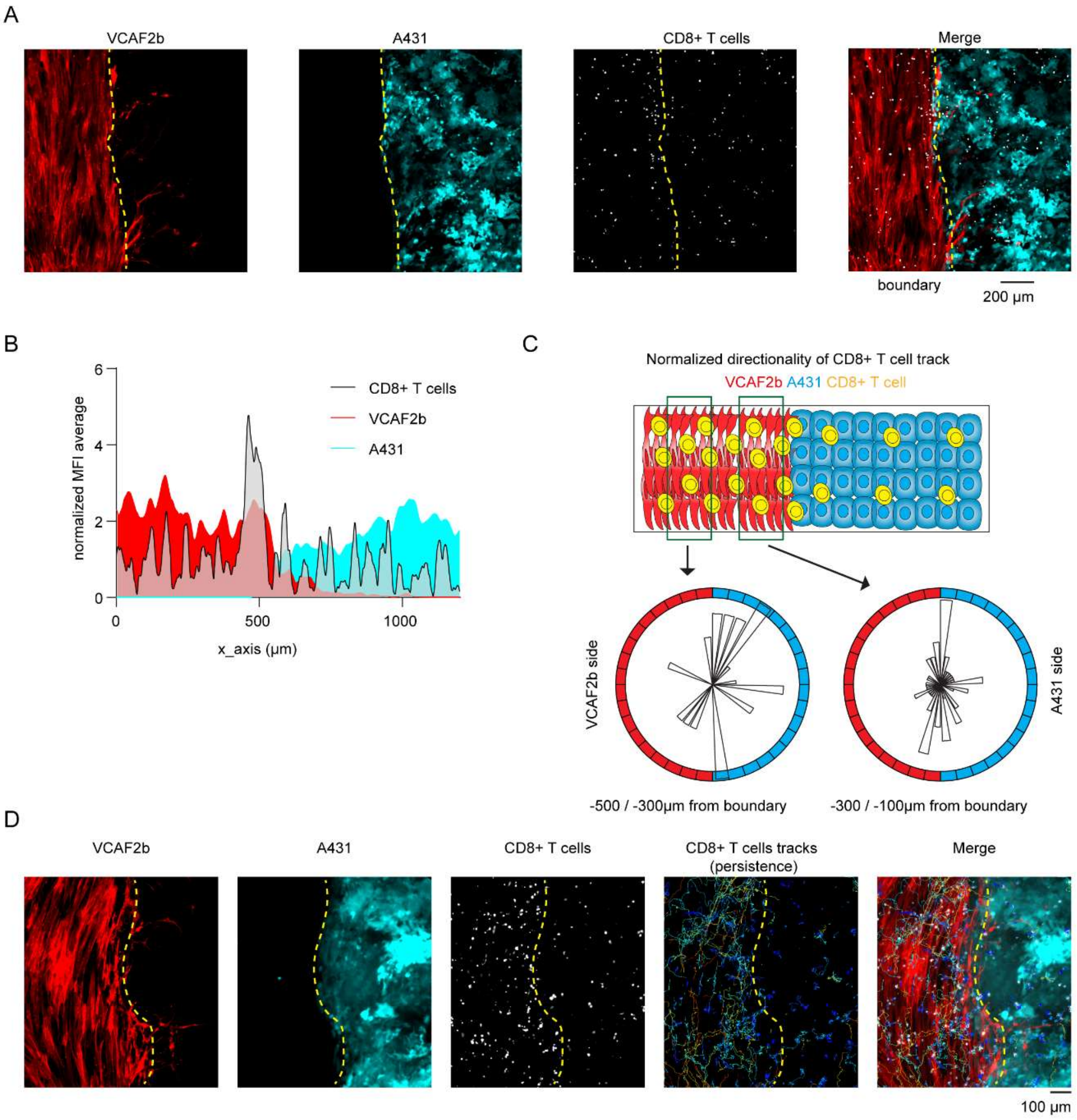

Supplementary Figure 7

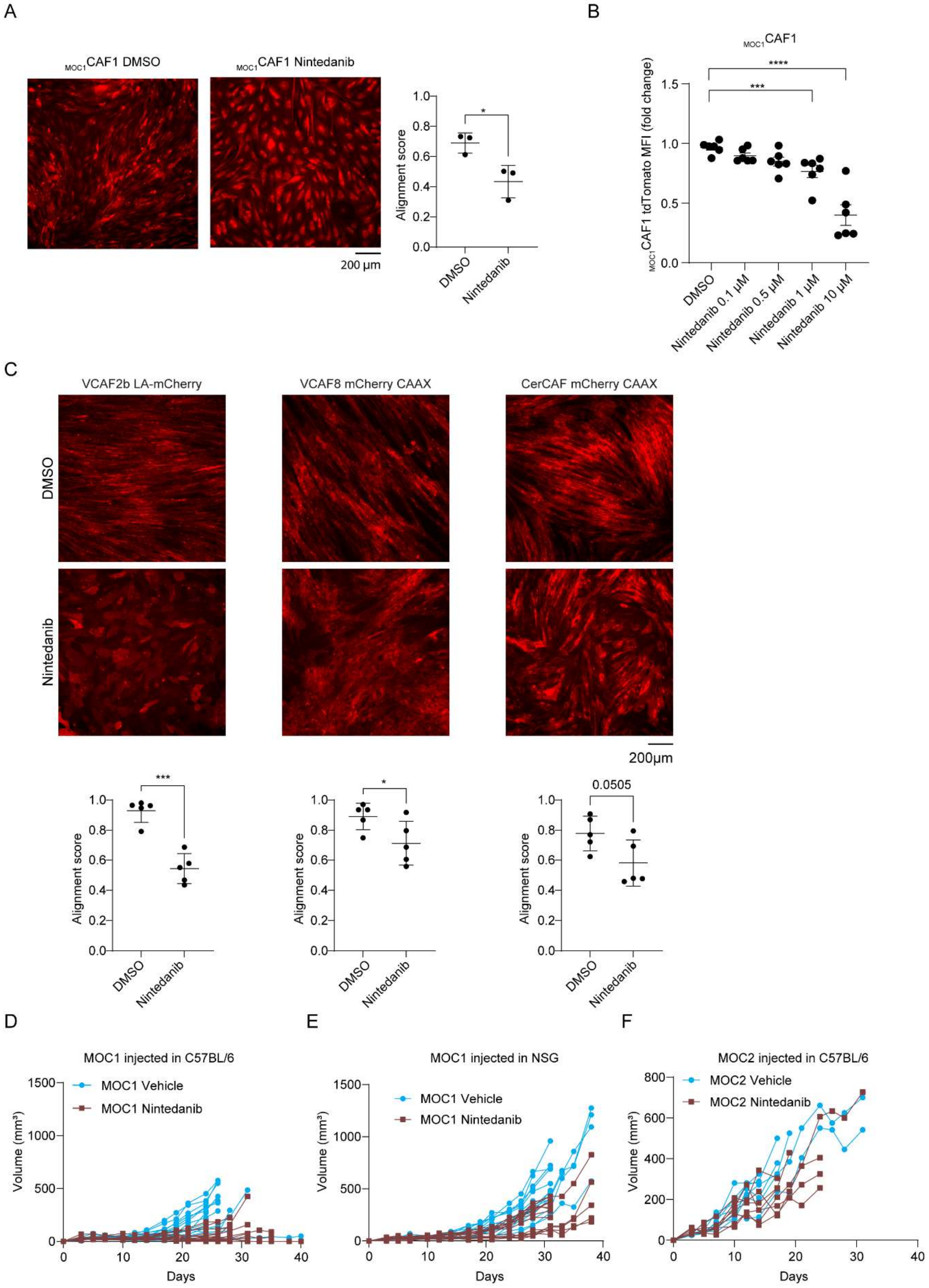

### Supplementary Figure 8

A

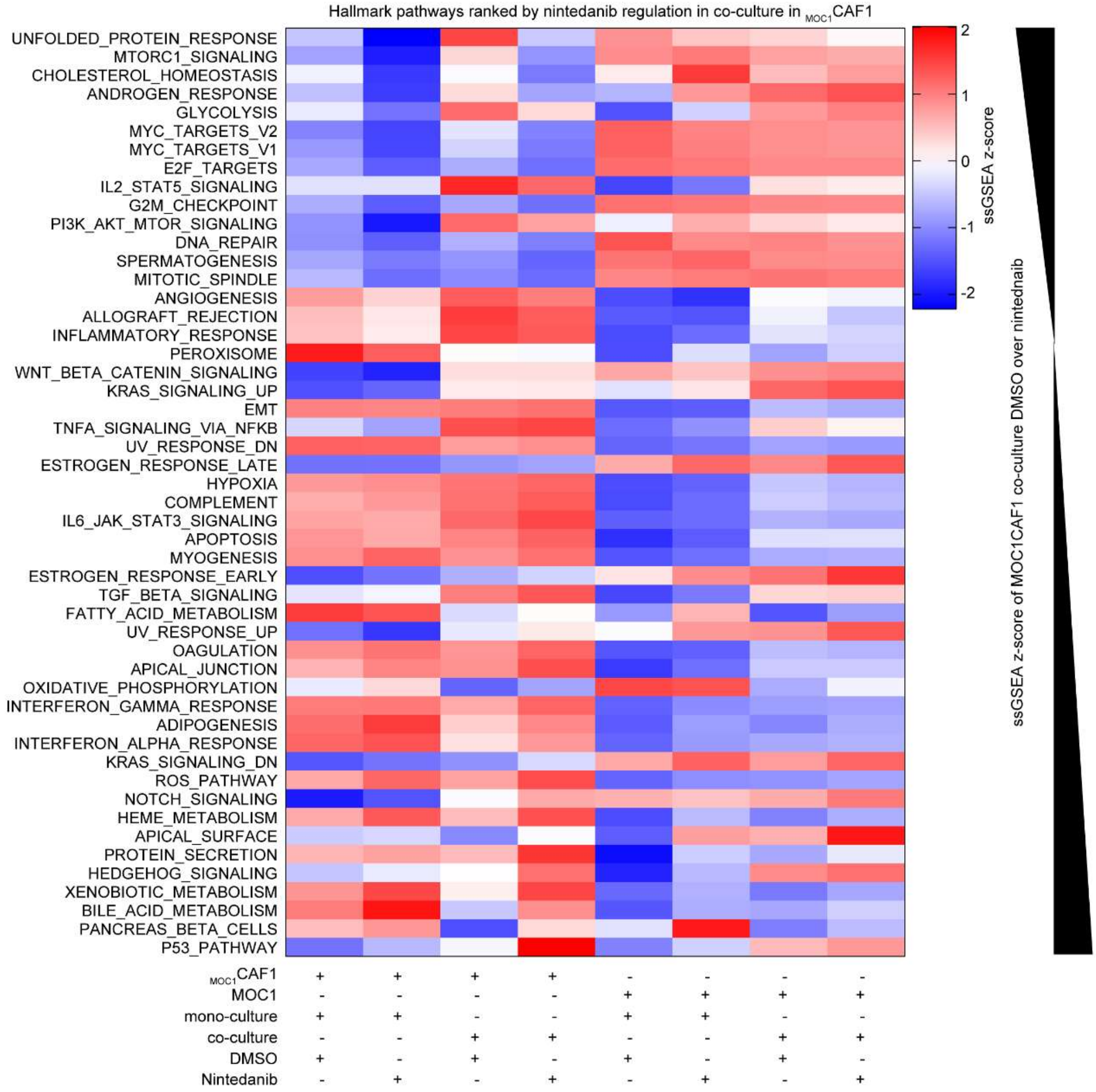


B

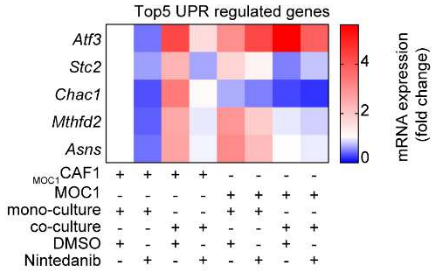

Supplementary Figure 9

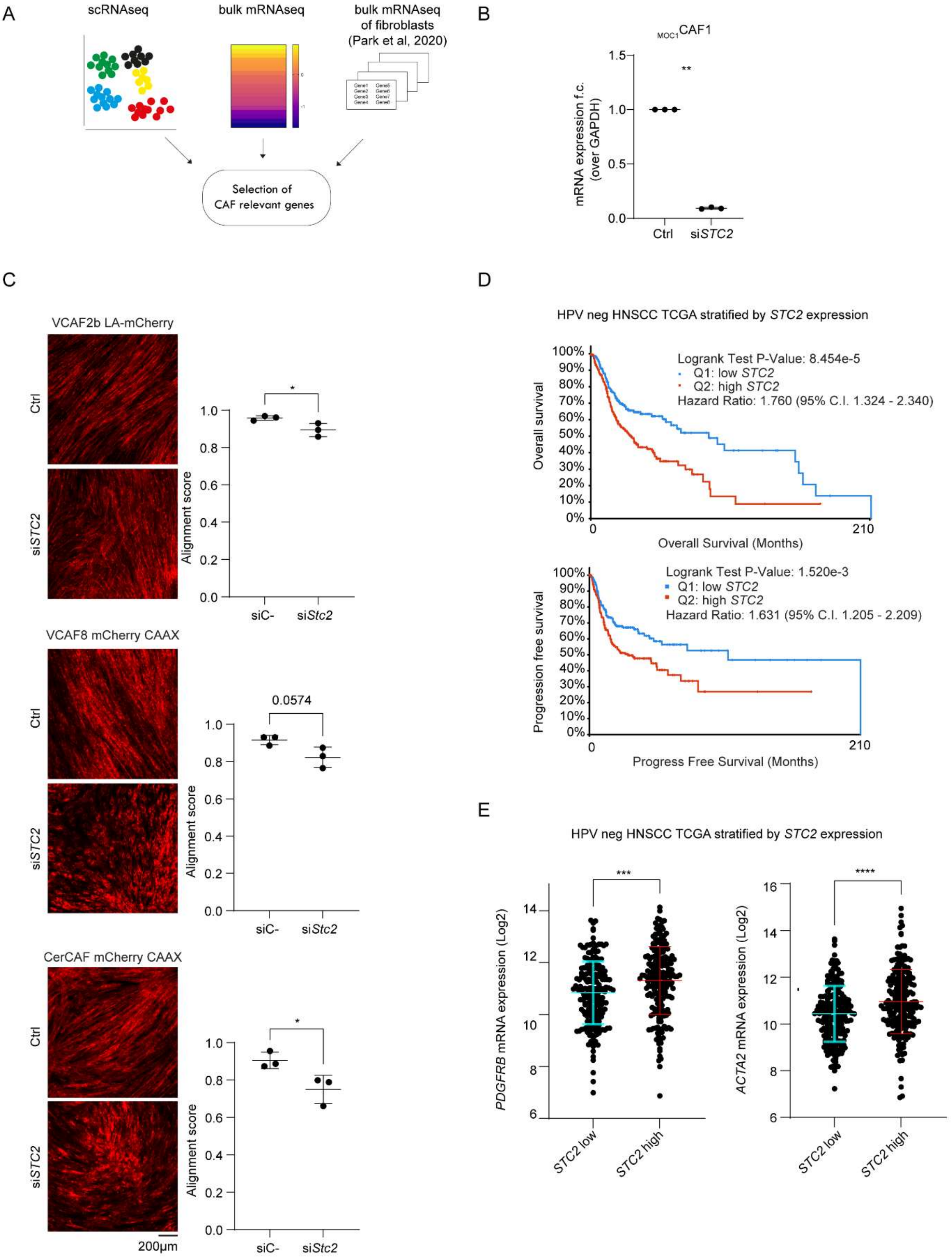

### Supplementary Figure 10

A

Descriptive analysis of $Ca^{2+}$ spikes

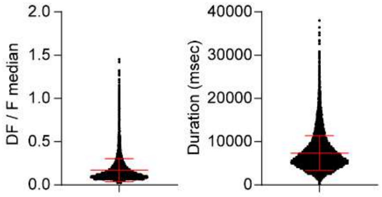


| | Delta F / F median | Duration (msec) |
|---|---|---|
| Minimum | 0.022 | 200 |
| 25% percentile | 0.090 | 4600 |
| 50% percentile | 0.127 | 6400 |
| 75% percentile | 0.200 | 9000 |
| Maximum | 1.453 | 38000 |
| Mean (95% C.I.) | 0.171 (0.169 - 0.173) | 7350 (7290 - 7410) |

B

Multicellular cluster spike counts

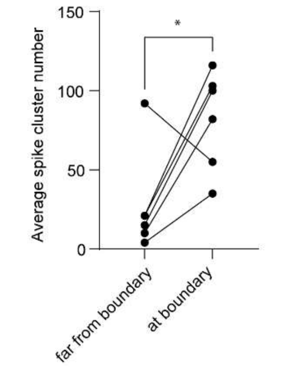


C

Cluster size

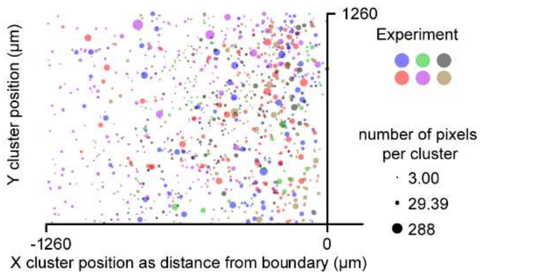


D

Multicellular cluster spike size

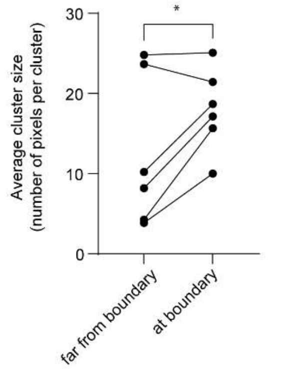


E

Cluster duration

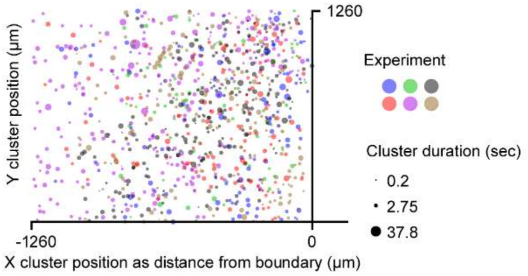


F

Multicellular cluster spike time duration

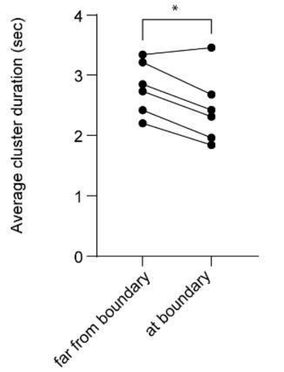
